# Reading ahead: Localized neural signatures of parafoveal word processing and skipping decisions

**DOI:** 10.1101/2025.04.25.650296

**Authors:** Graham Flick, Liina Pylkkänen

## Abstract

Visual reading proceeds fixation-by-fixation, with individual words recognized and integrated into evolving conceptual representations within only hundreds of milliseconds. This relies, in part, on interactions between cognitive and oculomotor systems, such that linguistic properties of words influence eye movements and fixation durations. When and where do these influences arise in the neural processing of an incoming word? To answer this, we combined magnetoencephalography (MEG) with eye-tracking in a natural story-reading paradigm. We replicated past findings that word frequency and predictability have additive influences on fixation durations. Next, we identified putative generators of these influences in localized brain activity time-locked to fixation onsets. Both properties independently influenced neural responses in left occipitotemporal and ventral temporal areas, at latencies early enough to influence subsequent saccade planning. These effects began in posterior areas (the left lingual gyrus, lateral occipital cortex) during parafoveal word processing, and shifted more anteriorly (the inferior temporal and parahippocampal gyri) when the word was fixated in foveal vision. Evidence for parallel processing of both parafoveal and foveal words was observed in the left posterior fusiform, which housed near-simultaneous effects of both the fixated and upcoming words’ frequency and surprisal. We also found that parafoveal processing in this region, together with the left middle temporal gyrus, distinguished whether an upcoming word was skipped or fixated. These results suggest that during natural visual reading, word recognition and integration begin parafoveally, underpinned by a left-lateralized occipitotemporal system, where word processing rapidly exerts downstream influences on eye movement decisions.

## 1. INTRODUCTION

The frequency and predictability of a given word are two of the strongest predictors of how long a visual reader’s eyes will linger on that word (Rayner, 1998; Sereno Rayner, & Posner, 1998; Sereno, Brewer, & O’Donnell, 2003). A central goal for the cognitive neuroscience of reading – and high-level vision more broadly – is to uncover how these influences are implemented mechanistically in the brain. Yet, despite great strides in neurobiological models of reading (e.g., Dehaene et al., 2010; Grainger, Dugau, & Ziegler, 2016), the fundamental question remains unanswered: How does neural activity give rise to linguistic influences on eye movements during visual reading?

Amongst past work that has examined word frequency and predictability, there is an often-discussed discrepancy related to fixation durations and brain responses (see Kretzschmar et al., 2015; Sereno et al., 2020; Huizeling et al., 2022). On one hand, eye-tracking studies have found additive effects of the two properties on fixation durations (e.g., Kliegl, Grabner, Rolfs, & Engbert, 2004; Staub, 2015). On the other, electroencephalography (EEG) studies have reported interactions between the properties (e.g., Van Petten & Kutas, 1990, 1991; Payne, Lee, & Federmeier, 2015). These interactions, which cannot be explained by paradigmatic differences (Kretzschmar et al., 2015), have commonly been observed at the N400 component (300-400 ms after word presentation; Kutas & Hillyard, 1980, 1984), where frequency effects attenuate in increasingly predictive contexts.

As the N400 peaks later than typical fixation durations (225-250 ms; Dimigen et al. 2011), and since the neural effects do not match the additive pattern, the field is without a mechanism that explains when and where, in brain activity, frequency and predictability assert their influences on eye movements. Addressing this issue requires recording neural responses to individual gaze fixations, with the spatiotemporal resolution to characterize discrete brain areas. In contrast, many previous studies relied on EEG (largely limited to descriptions at the scalp, e.g., Kreztschmar et al., 2009) or functional magnetic resonance imaging (fMRI, limited to blood flow changes across seconds, e.g., Henderson, Choi, Luke, & Desai, 2015; Schuster et al 2016), and/or studied brain responses in serial word presentation paradigms (e.g., Huizeling et al., 2022).

Here, we harnessed advances in co-registration methods (Nikolaev et al., 2016; Degno & Liversedge, 2020; Himmelstoss et al., 2020) in a simultaneous magnetoencephalography (MEG) and eye-tracking study of visual reading. The spatiotemporal resolution of MEG (Baillet, 2017), combined with eye-tracking and a deconvolution analysis (Ehinger & Dimigen, 2019), allowed us to relate localized brain activity to properties of fixated and parafoveal words, time-locked to fixation onsets. With this approach, we asked where and when word frequency and predictability influenced neural activity. Past studies using isolated word presentations have consistently implicated left occipitotemporal areas in bottom-up word recognition (Tarkiainen et al., 1998; Vinckier et al., 2007; Gwilliams, Lewis, & Marantz, 2016). Others found that contextual constraints influence orthographic processing, an early stage of word recognition (Laszlo & Federmeier, 2009; Caliskan et al., 2023), and demonstrated influences of these constraints in the same or nearby occipitotemporal areas (Dikker, Rabagliati, & Pylkkänen, 2009; Henderson, Choi, Lowder, & Ferreira, 2016; Schuster et al., 2021; Huizeling et al., 2022).

Based on these findings, we hypothesized that a word’s frequency and predictability – the latter operationalized as surprisal – would both influence activity elicited by fixations, in left occipitotemporal areas. Furthermore, we hypothesized that the effects would match the additive pattern observed in fixation durations and arise early enough to influence saccade planning, explaining the influences on eye movements. To facilitate this, and since recent work has suggested that both lexical and integration operations begin parafoveally (Pan, Popov, Frisson, & Jensen, 2023; Pan, Frisson, Federmeier, & Jensen, 2024), we expected that these effects would emerge in left occipitotemporal cortex prior to fixation. This would provide evidence for parallel visual processing in natural reading (Snell & Grainger, 2019) and identify the first port of call where word frequency and predictability exert their influences on eye movement behaviours.

## 2. MATERIALS AND METHODS

### 2.1 Participants

Thirty-two participants (22 women, 7 men, 3 undisclosed, mean age = 23.9 years, range = 18-55 years) took part in the study. This target sample size was defined based on those of past MEG studies of serial word reading (e.g., Gwilliams et al., 2016; Flick, Abdullah, & Pylkkänen, 2021) and similar sample sizes in co-registered reading studies using EEG or fMRI (Kretzschmar et al 2015; Henderson et al., 2015, respectively). All participants were fluent English speakers, with normal or corrected-to-normal vision and normal hearing, and without a history of linguistic or cognitive impairment. Eight further participants partially completed the procedure but had their data removed due to issues related to eye-tracking calibration or MEG data acquisition (n = 7) or a failure to follow instructions (n = 1). Of the thirty-two, 2 participants had their data removed before eye movement or MEG analysis due to issues in the eye-tracking data, and 2 participants were not included in the MEG analysis due to issues in source localization. This left a final sample of 28 participants in the MEG data analysis.

### 2.2 Experimental Design

Each participant read 216 short stories made of two sentences, selected from a subset of the total stimulus set, which consisted of 864 stories. Approximately one half of the complete set of stories were adapted from prior work by Sereno et al. (2020). This subset of the stimuli was modified to replace words typical of United Kingdom English with those more typical of American English. The remaining stimuli were generated following the structure of the set developed by Sereno and colleagues. Although they were not the focus of the present investigation, in the original design of Sereno et al., the second sentences in each story (the “target” sentence) contained a specified target word that fell into a low or high frequency category. Each target sentence occurred twice in the complete stimulus set used here: once with a context sentence that predicted the occurrence of the target word and once with a context sentence that did not, creating a manipulation of high versus low predictability. In the current study, stimuli were pseudo-randomly assigned to each participant such that they did not encounter any given target sentence in both of its contexts.

Rather than focusing on only the target words, the present investigation examined fixations on all words across the two sentences that made up each story. This increased the number of gaze fixations for analysis from an upper bound of 216 (one per story, per participant) to an upper bound defined by the total number of fixations per participant. For each word, we defined its length in number of letters, lexical frequency (logarithm-transformed frequency values extracted from the English Lexicon Project; Balota et al., 2007), and lexical surprisal, defined as the negative logarithm of the probability of each word, conditioned on those that occurred prior. Surprisal was estimated using the pre-trained GPT-2 large language model (Radford et al., 2019), selected because it has been shown to produce incremental surprisal estimates that correlate with human brain activity during language comprehension and word embeddings that map linearly onto human brain activations (Caucheteux, Gramfort, & King, 2022; Goldstein et al., 2022, Goldstein et al. 2024). We retrieved the pre-trained GPT-2 model from the *Hugging Face Transformers* Python Library (Wolf et al., 2020) and estimated surprisal using the *minicons* Python wrapper (Misra, 2022), with a window size that increased from 1 to a maximum of 12 context words, beginning on the second word in each story. For the eye-tracking data analysis, we also extracted each word’s letter bigram frequency from the English Lexicon Project.

### 2.3 Procedure

All participants provided informed consent prior to taking part in the experimental procedures, which took place in the MEG and MRI facilities of New York University. All procedures were approved by the Internal Review Board of New York University. Prior to MEG recordings, the shape of each participant’s head and the locations of three fiducial landmarks (nasion and tragi), as well as those of five head position indicator coils attached to the forehead and beside each tragus, were digitized in a three-dimensional model using a Polhemus FastSCAN system (Polhemus, Vermont, USA). Participants then entered the magnetically shielded room (MSR) that housed the MEG system, where they lay supine on a horizontal bed. Stimuli were back projected onto a display screen approximately 44 cm above the eyes. The viewing distance varied approximately 1-5 cm depending upon the size of the participant’s head.

Participants were instructed that they would be reading short stories made of two sentences. The initial sentence in each story was preceded by a green fixation cross that marked the position of the first letter in the first word, approximately 3 cm from the left boundary of the screen and 2 cm above the middle vertical position. After a delay of 300 ms, the first sentence in the story replaced the green fixation cross. Four characters subtended a horizontal angle of approximately 1.5°. All sentences were displayed in the Courier New font. Participants read the first sentence in the story naturally with eye movements and pressed a button in their left hand when they were finished. A grey fixation cross then appeared for 300 ms, again marking the starting position of the first letter, followed by the second sentence in the story. Participants read this naturally with eye movements before pressing the button to indicate that they were finished.

On one third of trials, a second grey fixation cross appeared after participants read the second sentence in the story. If this occurred, participants read a third sentence then responded to a yes/no question asking if the continuation of the story was natural or unnatural. Half of the continuation sentences were written to be natural fits with the story they appeared in, whereas the other half were randomly permuted continuations intended for other stories, from the set of stimuli that the participant would not read. These continuations were inserted into the trials to motivate participants to pay attention and read each story carefully. Because the random permutation of continuations often resulted in unnatural story endings that participants nevertheless interpreted as sensible, there was not an unambiguously correct answer on every trial. For this reason, we refrained from analyzing the behavioural accuracy data for these judgments. Fixations and MEG data during the reading of the third sentences were not included in any of the analyses.

Participants completed a brief practice of the reading task in the magnetically shielded room prior to beginning the experiment. They then completed six blocks of the task while MEG and eye-tracking data were simultaneously recorded. Each block contained thirty-six stories and took approximately 10 minutes to complete, including eye-tracker calibration and recording setup. Participants read 532 analyzed sentences in total. Following completion of the MEG procedures, participants were asked to return to have a T1 anatomical MRI scan. MRIs were collected on a Siemens 3T Prisma system. Seventeen participants completed the MRI session.

### 2.4 Data Collection & Co-registration

Continuous MEG data were recorded using a 156-channel Kanazawa Institute of Technology axial gradiometer system, with a sampling rate of 1000 Hz and online low- and high-pass filters of 200 Hz and 1 Hz, respectively. The 1 Hz high-pass filter was necessary due to external electromagnetic interference in the low frequency range, generated by the surrounding New York City environment.

Eye-tracking data were collected with an SR Research Eyelink 1000 long-range system (SR Research, Mississauga, Ontario) with a 1000 Hz sampling rate. The eye-tracker was mounted above the participant as they lay supine. For each participant, we determined which eye was best captured by the tracker while the participant was positioned in the MEG, and recorded data from that eye. Calibration and validation routines were performed prior to each block of the reading task. For the first 12 participants, a nine-point calibration was performed, placing dots at each of the screen’s corners and mid-way between the corners. The resulting rectangle spanned 88% of the screen horizontally and 83% of the screen vertically. During these initial participants, it was regularly found that calibration points in the diagonal corners resulted in poor validation, with considerable time spent adjusting the positioning of the tracker. Thus, for the remaining participants, a five-point routine was used to reduce calibration time. Points were placed at cardinal directions of the original rectangle. Even with the reduced calibration grid, the routine captured the entire span of the sentence stimuli. Calibrations were accepted if the average error on the subsequent validation run met the SR Research standard definitions of “Good” or “Fair”, defined as an average error of less than 1 degree (worst error point less than 1.5 degrees) or an average error between 1 and 1.5 degrees (worst point error less than 2 degrees), respectively. Five blocks (approximately 3% of the dataset) in which calibration did not reach this level of accuracy were excluded from data analysis.

To co-register the MEG and eye-tracking data, the experiment control computer coordinated presentation of the stimuli to the participant’s screen using Pyschtoolbox (Brainard, 1997) in MATLAB (Mathworks, Natick, USA) and sent event codes to both recordings. A simultaneous trigger pulse was sent to the MEG system and a message to the eye-tracker acquisition computer, via ethernet connection, when a stimulus appeared on the screen. Delays between the sending of the visual stimulus and its appearance on the screen were adjusted using a photodiode, which recorded stimulus onsets directly in the MEG data. The timing of gaze fixations from the eye-tracking data were then used to define epochs of fixation-related fields in the MEG data, using the relative time of the fixation onsets from the onset times of the sentences, which were coded in both data modalities. During stimulus creation, the position of each word in each sentence was defined in a bounding box, including one half letter unit of white space on either side, and a buffer of 25 pixels above and below the word vertically. In a subset of participants whose eye-tracking data showed vertical drift, resulting in fixations above the line of text but not beyond its horizontal borders, we expanded the vertical range of the bounding boxes to capture these fixations. Fixations, saccades, and blinks were identified in the eye-tracking data using the standard Eyelink parser from SR Research. Using the positions of gaze fixations in the eye-tracking data, we related each epoch of MEG data to an individual word that was fixated at the instant in time.

### 2.5 Statistical Analysis

#### 2.5.1 Eye-tracking Analyses

An overview of the co-registration and data analysis is shown in Figure 1. We focused our analysis on first-pass first fixation durations, defined as the duration of time a participant viewed each word the first time they fixated on it, prior to advancing any further in the sentence.

**Figure 1.**
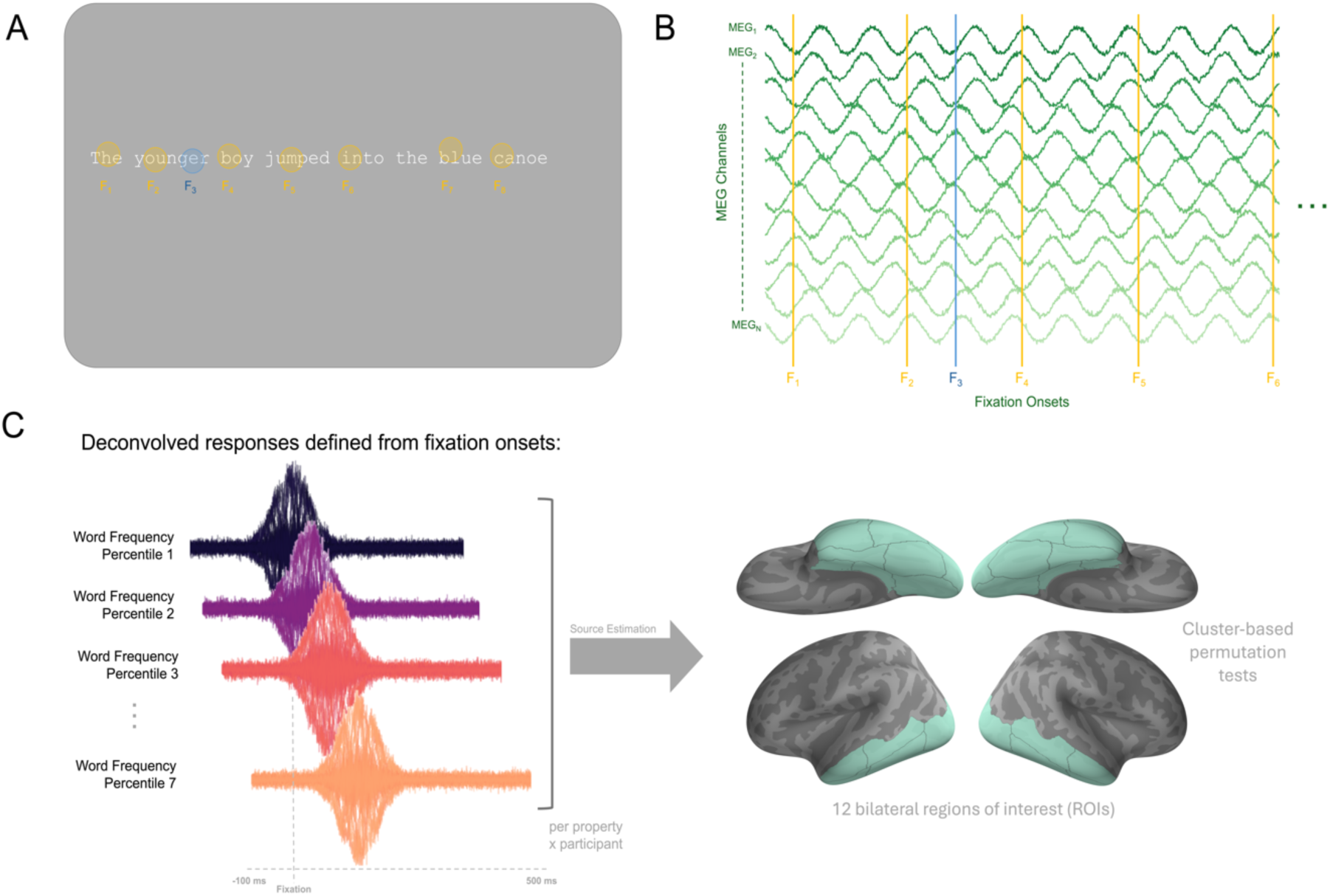
(A) Participants read short stories made of two sentences, presented one-at-time, while co-registered MEG and eye-tracking data were recorded. In this example, first fixations are shown in gold (e.g., F_1_, F_2_). A single re-fixation (F_3_) is shown in blue. (B) Fixation onset times were defined in the MEG data using the relative time from the start of each trial. (C) Deconvolution was used to generate estimated responses from each MEG channel, time-locked to the onset of first-pass first fixations. We modelled responses to four properties of interest (fixated word frequency and surprisal, right neighbouring word frequency and surprisal) at seven percentiles from each participant’s data. The estimated fixation-related were localized to the cortical surface, where cluster-based permutation tests were used to assess source activity in 12 bilateral regions of interest (ROIs; C, right).

Fixation durations were summarized with descriptive statistics and analyzed with linear mixed effects models in R (R Core Team, 2022) using the lme4 package (Bates et al., 2015). Likelihood ratio tests were used to assess the contribution of each predictor variable (Winter, 2013), by comparing models with and without that predictor. Following past work using co-registered EEG and eye-tracking (Ehinger & Dimigen, 2019), we focused the analysis on open class words (nouns, adjectives, verbs, and adverbs) and excluded fixations that were less than 50 ms, greater than 750 ms, and those that occurred within 500 ms of a preceding blink. All models included by-participant random intercepts, but not by-participant random slopes due to convergence issues.

#### 2.5.2 MEG Pre-processing

MEG data were preprocessed using the MNE-Python (Gramfort et al., 2013, 2014) and Eelbrain (v.037, Brodbeck et al., 2023) libraries in the Python computing environment. Data from each participant were first cleaned of externally generated electromagnetic noise using the Continuously Adjusted Least Squares Method (Adachi et al., 2001). The continuous data were low-pass filtered at 40 Hz and visually inspected to identify sections of residual noise (e.g., muscle contractions or other movements) in the data, which were annotated as bad segments and dropped from subsequent analysis. Independent Components Analysis (ICA) was used to decompose the continuous MEG data into latent components that explained 95% of the variance in the dataset. For each participant, we identified the component(s) that showed the sensor topography characteristic of a saccade artifact: a uniform field at MEG channels located above the eyes (see Hari & Puce, 2023, for detailed discussion of electrophysiological ocular artifacts). We validated this interpretation of the saccade-related components by using the same ICA solution to later decompose each participant’s fixation-related epochs of MEG data, confirming that the time course of component saliences showed highest magnitudes surrounding the onset and offset of the saccade. Saccade-related components, as well as components related to well-known external noise sources, cardiac activity, and blinks, were removed from the continuous data before further analysis.

#### 2.5.3 Deconvolution & Source Localization

Past work has demonstrated that the rapid nature of eye movements results in neural responses that overlap in time and which, if not properly disentangled, can lead to spurious differences in estimated activity related to an event of interest (a problem not unique to eye-tracking paradigms; Ehinger & Dimigen, 2019). To unmix the neural responses, we adopted a deconvolution analysis using the Unfold MATLAB toolbox (Ehinger & Dimigen, 2019). This approach involves solving for a vector of parameter weights, or beta coefficients, corresponding to b in the general linear model y = Xb + e, where y is the continuous MEG data from one channel, e is a vector of residuals, and X is a *time-expanded* design matrix that defines the local window of interest around the onset of target events, throughout the duration of the recording (see Ehinger & Dimigen 2019 for details). The resulting beta coefficients span that local time-window around the onset of each event (i.e., −100 to 500 ms around the onset of first fixations) and capture the influence of a modelled property on neural activity in that window, while reducing the influence of modeled events that occurred before or after it, such as the onset of the sentence stimulus, and previous or upcoming saccades and fixations. The deconvolution method has been widely used in analyses of neural time series data (e.g. Coco et al., 2020; Norman et al., 2021) and similar approaches for continuously varying stimuli (e.g., audio narratives) have become increasingly popular in M/EEG research (i.e., temporal response functions: Cross et al., 2016; Brodbeck et al., 2023)

We defined a design matrix with nuisance columns indicating the onset of each sentence on the screen, the onsets of fixations on closed class words, and the onset of any refixations or fixations not on words, during the reading of that sentence. Each of these events was modelled using only an intercept term in the −100 to 500 ms window. We then added columns capturing the primary events of interest: the onset of first-pass first fixations on open-class words, with an analysis window of −100 to 500 ms around those onsets. Responses to these events were modeled with an intercept term and parametric predictors: a categorical variable capturing the length of the fixated word (short: 3 letters or less; medium: 4 to 7 letters, inclusive; long: more than 7 letters), and continuous predictors capturing the amplitude of the incoming saccade, the lexical frequency of the fixated word, the surprisal of the fixated word estimated from GPT-2, and the length (short, medium, long), frequency, and surprisal of the right neighboring word. Continuous predictors were modelled using spline regression (Wood, 2017), with seven splines defined across quantiles of the predictor’s range. Models were fit for each participant before the resulting beta coefficients were used to generate predicted responses from seven percentiles of each modelled continuous property, marginalizing over all other predictors in the model. This yielded estimates of the effect of each predictor, holding all others constant, and disentangled in time from events that preceded or followed.

The predicted magnetic flux responses – one for each level of each predictor, for each participant – took the form of sensor-by-time data matrices that spanned 100 ms before fixation onset to 500 ms afterward. To characterize the neural generators of these responses, we used noise-normalized minimum-norm estimation (Dale et al., 2000) to convert each data matrix to a time course of predicted source activity. T1 anatomical MRIs were used to define cortical surface models via Freesurfer’s automated segmentation algorithms (recon-all; Dale, Fischl, & Sereno, 1999; Desikan et al., 2006; Fischl, Sereno, & Dale, 1999; Fischl et al., 2004). The MEG data from each participant were co-registered to their anatomical surface based on the digitized locations of the fiducial points and the shape of the scalp. For those participants who did not have an anatomical MRI, freesurfer’s *fsaverage* template brain was scaled and stretched to match the location of the participant’s fiducial points and the shape of their scalp. Channel noise covariance matrices were estimated from a three-minute empty-room recording collected just prior to or following the participant’s experimental session, using the automated model selection and regularization method of Engemann & Gramfort (2015). Source estimation parameters were defined based on past work that has examined the sensitivity of MEG recordings to stages of visual word recognition that occur rapidly after foveal word presentation (Gwilliams et al., 2016; Flick et al., 2021). In keeping with those past studies, source estimates were generated using “fixed” orientation, constrained based on the geometry of the cortical surface, with cortical patch statistics used to define the orientation normal to the surface (Lin et al., 2006).

#### 2.5.4 Region of Interest Analyses

Once localized, predicted responses were analyzed in regions of interest (ROIs) using spatiotemporal cluster permutation tests (Maris & Oostenveld, 2007) with 5000 permutations to define surrogate null distributions. ROIs were defined in the cortical surface using the automated Desikan-Killiany parcellation (Desikan et al., 2006). To provide a comprehensive examination of areas implicated in visual word recognition, we included a bilateral set of regions that spanned early visual cortex (cuneus, lingual gyrus, pericalcarine cortex), occipitotemporal cortex (lateral occipital cortex, fusiform gyrus, inferior temporal gyrus) and the lateral and medial temporal lobes (entorhinal cortex, parahippocampal cortex, middle temporal gyrus). We excluded inferior and superior frontal areas due to the lower spatial accuracy of MEG source localization in these anterior sites (Hauk et al., 2011), which may be intensified by the supine position of participants in the MEG system, where the head was pressed against the back of the system’s helmet. To provide greater spatial insight, the fusiform, inferior temporal, and middle temporal gyri were split into posterior and anterior sections, defined as an even split along the range of their posterior-to-anterior span.

For each ROI, clusters were formed from F-statistics calculated across the estimated responses of each level of each predictor, using a one-way ANOVA. Clusters were defined using cluster-free threshold enhancement (Smith & Nichols, 2009). Altogether, these methods identified areas where there was significant variability, above that expected by chance alone, in neural activity related to each of the predictors of interest. Primary analyses were performed on localized neural activity from 0 to 225 ms following fixation onset. The time window was motivated by the typical average duration of fixations reported in past literature, which closely matched that found here (see Section 3.1), and was intended to capture neural activity prior to the average onset of the following saccade. Moreover, by using a deconvolution analysis that considered the onsets of all fixations, we dis-entangled neural responses to subsequent eye movements from those under analysis in this time window, such that effects were associated with the onset of the currently examined fixation, rather than any that followed (Ehinger & Dimigen, 2019). This also enabled us to examine effects of the upcoming, right neighbouring word’s properties before it was fixated, in the same 0-225 ms time window relative to the fixation on the preceding word.

All clusters containing a minimum of three sources and a duration of at least 10 ms were assessed for significance. Cluster p-values were corrected for multiple comparisons using the family-wise error rate accounting for arbitrary dependencies (Benjamini & Yekutieli, 2001).

Source analyses examining fixated and parafoveal word frequency and surprisal were corrected by taking into consideration all tests across predictors and ROIs, unless stated otherwise below. Exploratory tests on later activity, from 225-500 ms, were subjected to the same multiple comparison correction as the original set conducted from 0-225 ms, but not for the total number of comparisons across both analyses. A corrected threshold of p < 0.05 was considered statistically significant.

We note that the polarity of estimated source amplitudes can depend on several factors, including the location in the cortical folding and distance from the MEG system sensors, which vary across participants. For these reasons, we refrain from interpreting differences in response magnitudes as relative activations or deactivations. Instead, the presence of significant effects is interpreted as evidence that the property influences activity within the examined ROI and analysis time window. For visualization, cluster time courses were extracted by finding the dominant direction or orientation of all sources in that area and applying a sign-flip to those that had the opposite polarity.

## 3. RESULTS

### 3.1 Frequency and surprisal additively influence first-pass first fixation durations

The mean first-pass first fixation duration on open class words was 225.05 ms (SD = 92.75 ms), closely matching that reported in past studies, including previous co-registration paradigms (e.g., Rayner, 1978; Choi, Desai, & Henderson, 2014). To examine the impact of each predictor, we compared a series of nested linear mixed effects models using likelihood ratio tests.

We began by comparing a model that attempted to explain first-pass first fixation durations using three nuisance predictors (word length, mean letter bigram frequency, and ordinal position in the sentence) to models that additionally contained word frequency or surprisal. Adding either frequency (X^2^(1) = 41.01, p < 0.001) or surprisal (X^2^(1) = 28.13, p < 0.001) resulted in significant model improvement. The same significant improvements were found when adding either predictor to models that contained the other, as well as the original nuisance predictors (adding frequency: X^2^(1) = 19.52, p < 0.001; adding surprisal: X^2^(1) = 6.637, p = 0.009).

In all models, coefficients on the fixed effect of frequency were negative, demonstrating shorter fixation durations on higher frequency words (−1.582 +/-0.358 [standard error], extracted from the full model), whereas coefficients on fixed effect of surprisal were positive, demonstrating greater fixation durations on more surprising words (0.592 +/-0.230). Finally, adding the interaction between word frequency and surprisal did not improve model performance (X^2^(1) = 1.686, p = 0.194). Altogether, these results match previous findings that the two predictors display additive, rather than multiplicative, influences on first fixation durations (e.g., Kennedy et al., 2013; Kretzschmar et al., 2015; Staub, 2015).

### 3.2 Frequency and surprisal both influence neural activity in left occipitotemporal ROIs

Next, we assessed how fixated word frequency and surprisal each influenced neural activity from 0 to 225 ms after first-pass first fixation onsets. Deconvolved neural responses to seven levels of each predictor’s magnitude were localized to the cortical surface and subjected to spatiotemporal cluster-based permutation tests to identify ROIs that showed impacts of each property. As shown in Figure 2 (A & B), significant effects were found throughout occipitotemporal, lateral and medial temporal, and early visual ROIs. Time-courses from significant clusters in the left parahippocampal cortex, anterior inferior temporal gyrus, and posterior and anterior fusiform gyrus are shown in Figure 3. Cluster test results are summarized in Supplementary Tables 1 and 2.

**Figure 2.**
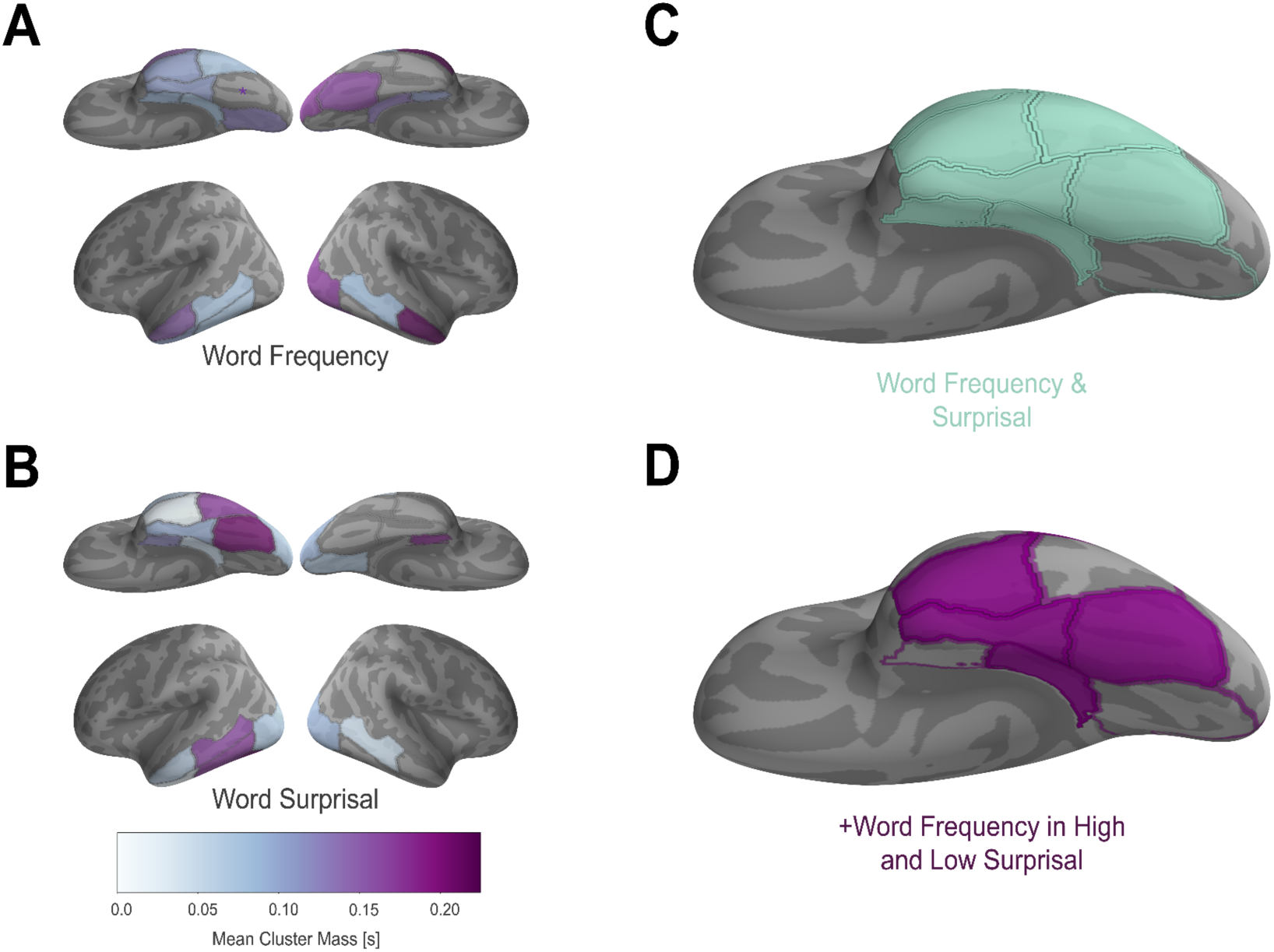
Effects of word frequency and surprisal on localized neural activity from 0-225 ms after the onset of first-pass first fixations. (A) ROIs displaying significant effects of the fixated word’s frequency are shaded. The left posterior fusiform gyrus is not shaded but marked with an asterisk as frequency effects did not survive correction for multiple comparison. Mean cluster temporal mass was calculated as the average of all time points in clusters within each ROI. (B) ROIs displaying significant effects of the fixated word’s surprisal. (C) The overlap of ROIs that showed both frequency and surprisal effects of the fixated word on the ventral surface of the left hemisphere, including the left posterior fusiform. (D) Those ROIs in (C) that additionally showed effects of frequency across both high and low surprisal contexts. Cluster test details are listed in Supplementary Tables 1 and 2.

**Figure 3.**
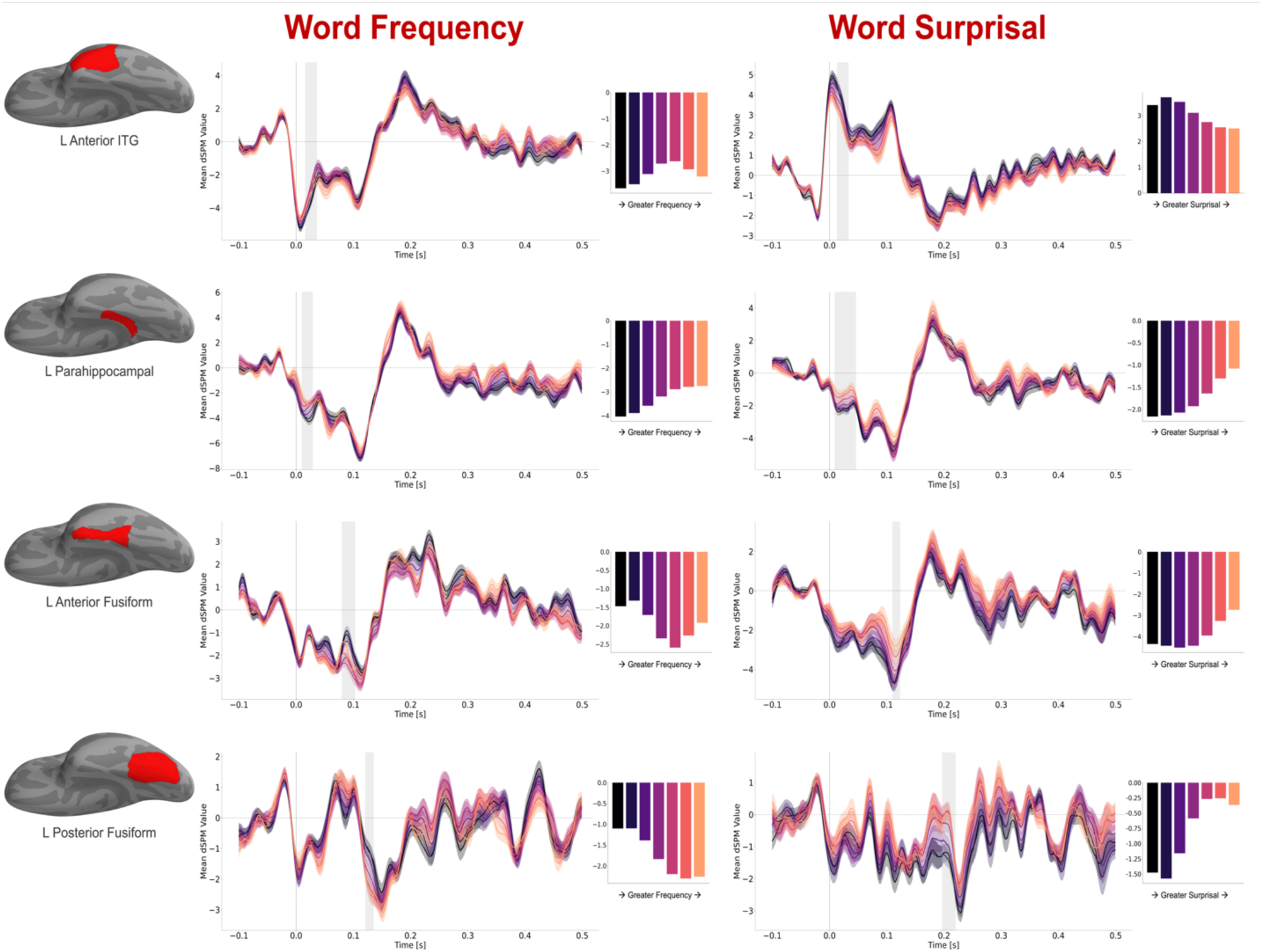
Frequency (left column) and surprisal (right column) effects on post-fixation responses. Top-row: Cluster time courses extracted from the left anterior inferior temporal gyrus ROI, showing notably early effects of frequency (16-36 ms) and surprisal (13-33 ms). Second row: Effects of frequency (10-29 ms) and surprisal (9-46 ms) in the left parahippocampal ROI. Third row: Frequency (80-103 ms) and surprisal (110-123 ms) effects in the left anterior fusiform ROI. Bottom row: Frequency (121-137 ms) and surprisal (197-220 ms) effects in the left posterior fusiform ROI. Shaded windows of each time course indicate the temporal extent of each cluster. Bar plots to the right indicate the mean estimated activity extracted from the cluster for each level of the predictor.

Fixated word frequency effects were found within 225 ms after fixation onset bilaterally in the pericalcarine cortex (corrected p-values = 0.019 in both hemispheres), parahippocampal cortex (left p = 0.019, right p = 0.020), entorhinal cortex (left: p = 0.019; right: p = 0.022), and the middle temporal gyrus (left: p = 0.019 anterior, p = 0.024 posterior; right: p = 0.024 anterior, p = 0.039 posterior). Left lateralized effects of frequency were found in the lingual gyrus (p = 0.019), anterior fusiform gyrus (p = 0.019), and inferior temporal gyrus (both anterior, p = 0.020, and posterior, p = 0.019; note that a corrected p-value of 0.019 indicates a cluster that was larger than any observed in the permutation distribution). An effect of frequency was also observed in the left posterior fusiform but did not survive correction for multiple comparisons (uncorrected p-value = 0.009; corrected = 0.094). Additional right hemisphere effects were observed in the lateral occipital cortex (p = 0.019) and the posterior fusiform gyrus (p = 0.019).

Many of the observed clusters showed effects of frequency (and surprisal) with strikingly early latencies, such as 30 ms after fixation onset. We return to this issue in Section 3.3, when discussing the impact of parafoveal processing, which may facilitate these early effects.

Surprisal effects, shown in Figures 2(B) and 3, were observed in many of the same ROIs that displayed influences of frequency, with an apparent left-lateralization of overlap. Left hemisphere regions that showed both effects were the posterior and anterior sections of the left inferior temporal gyrus (both corrected p-values = 0.019), anterior fusiform gyrus (p = 0.019), and middle temporal gyrus (p = 0.019, anterior and posterior), the parahippocampal cortex (p = 0.019) and the entorhinal cortex (p = 0.019). Overlap was also found in the right lateral occipital cortex (p = 0.019) and the right entorhinal cortex (p = 0.019). Surprisal effects were additionally observed in the left posterior fusiform (p = 0.019, where frequency effects did not survive correction), left lateral occipital cortex (p = 0.030), the right lingual gyrus (p = 0.019), and the right cuneus (p = 0.028). Figure 2C displays the set of left ventral areas that showed both frequency and surprisal effects of the fixated word.

These results highlighted a predominantly left lateralized set of areas that housed effects of a fixated word’s frequency and surprisal, beginning rapidly after the eyes were placed upon that word. This suggests these areas show local additive influences of the two properties, similar to first fixation durations. Motivated by the degree of overlap that emerged, we further evaluated whether these regions contained effects of frequency independent of surprisal by following the approach of previous studies (Van Petten & Kutas, 1990, 1991; Dambacher et al., 2006) and testing for effects of word frequency in contexts of different predictability. For each participant, we defined fixations on words below and above the 35th and 65th percentiles of surprisal values, respectively, defined from all words viewed on first-pass first fixations, from that participant’s data. The resulting low and high surprisal events corresponded to predictive and non-predictive contexts. We considered ROIs that showed main effects of frequency and surprisal across all examined fixations, and robust effects of frequency in both high and low surprisal subsets, to provide the strongest evidence for locally independent influences of the two properties.

Effects of word frequency in both levels of surprisal were almost exclusively found in left hemisphere ROIs, and specifically in ventral temporal and occipitotemporal areas (Figure 2D, Supplementary Table 3). These effects were observed in both anterior and posterior sections of the left fusiform gyrus (anterior: high surprisal p = 0.019, low surprisal p = 0.037; posterior: high surprisal p = 0.019, low surprisal p = 0.034), left anterior inferior temporal gyrus (high surprisal p = 0.019, low surprisal p = 0.020), left posterior middle temporal gyrus (high surprisal p = 0.019, low surprisal p = 0.022), left parahippocampal gyrus (high surprisal p = 0.019, low surprisal p = 0.019), and left lateral occipital cortex (high surprisal p = 0.019, low surprisal p = 0.019). The right lateral occipital cortex also showed frequency effects in both levels of surprisal and was the only right hemisphere ROI to do so (high surprisal p = 0.019, low surprisal p = 0.032).

As reported above, main effects of surprisal were observed in all of these ROIs and main effects of word frequency, across all examined fixations, were observed in the left fusiform (posterior and anterior), anterior inferior temporal gyrus, and parahippocampal cortex (Figure 2D). These findings thus suggest independent influences of frequency and predictability on neural activity, predominantly in left ventral temporal and occipitotemporal areas, which match the observed influences of these properties on first-pass first fixation durations (see Section 3.1).

### 3.3 No evidence that frequency and predictability overlap at individual sources and time points

The previously described effects of frequency and surprisal were found in spatiotemporal clusters, within the encompassing time window of 0-225 ms post-fixation onset, and within the spatial boundaries of each ROI. In one region, the impact of surprisal could influence responses at a set of sources and/or time points that do not overlap with the effect of frequency (and vice versa). Another possibility is that within each ROI, the two properties influenced neural activity localized to the same set of constituent sources, and at the same time points, after fixation onset. This degree of co-localization would provide the strongest evidence that the two properties affect a shared stage of processing in visual word recognition.

To test this possibility, we extracted the frequency clusters and performed one-way repeated-measures ANOVAs across levels of word surprisal on mean activity localized to each cluster’s spatiotemporal extent. This would identify clusters that showed significant modulation by word surprisal, as well as word frequency. Of the forty-two frequency clusters identified across all ROIs, zero showed significant effects of word surprisal (all corrected p-values > 0.05). Likewise, of the thirty-seven total surprisal clusters identified from the original analysis, zero showed significant effects of word frequency within their spatiotemporal extents (all corrected p-values > 0.05). These results thus fail to find evidence for the strongest form of overlap between frequency and surprisal effects, as the influences do not appear to localize to common spatiotemporal windows.

### 3.4 Parafoveal influences of the right neighbouring word’s frequency and surprisal

How early do the properties of a word begin to influence neural activity? To address this question, we investigated the impact of a word’s frequency and surprisal before the eyes moved to view that word. Specifically, we examined the influence of the right neighbouring word’s properties on deconvolved neural activity from 0-225 ms, after the onset of first-pass first fixations on open-class words. Notably, this approach differs from examining the parafoveal processing of the open-class words themselves, prior to fixation upon them. We elected to take this approach as it allows a common set of fixations to be used in both analyses, thus paralleling the influence of the fixated words when considering parafoveal processing. It also removes complications that arise from time-locking to the offset of the previous saccade (i.e., looking backward from fixation onset) and the possibility of skipped words appearing before open-class words. Reciprocally, we were able to examine the differential processing of the subsequent words when they were skipped or fixated (see Section 3.6).

Shown in Figure 4, significant influences of the right neighbor properties were observed in many regions that showed effects of the same properties of the foveal word. Fixated and neighbouring word frequency both influenced activity in the left fusiform gyrus (anterior; p = 0.020, posterior: p = 0.019), lingual gyrus (p = 0.019), pericalcarine cortex (p = 0.019), parahippocampal cortex (p = 0.019), and entorhinal cortex (p = 0.024), as well as the right lateral occipital cortex (p = 0.019) and right posterior fusiform (p = 0.041). Neighbouring frequency effects were also observed in the left lateral occipital cortex (p = 0.019) and the right posterior inferior temporal gyrus (p = 0.049). Neighbouring and fixated word surprisal both influenced activity in the left fusiform gyrus (posterior: p = 0.019), posterior inferior temporal gyrus (p = 0.019), anterior middle temporal gyrus (p = 0.019), lateral occipital cortex (p = 0.019, and the right cuneus (p = 0.019) and right entorhinal cortex (p = 0.032). For details on the effects of the right neighbour properties, see Supplementary Tables 4 and 5.

**Figure 4.**
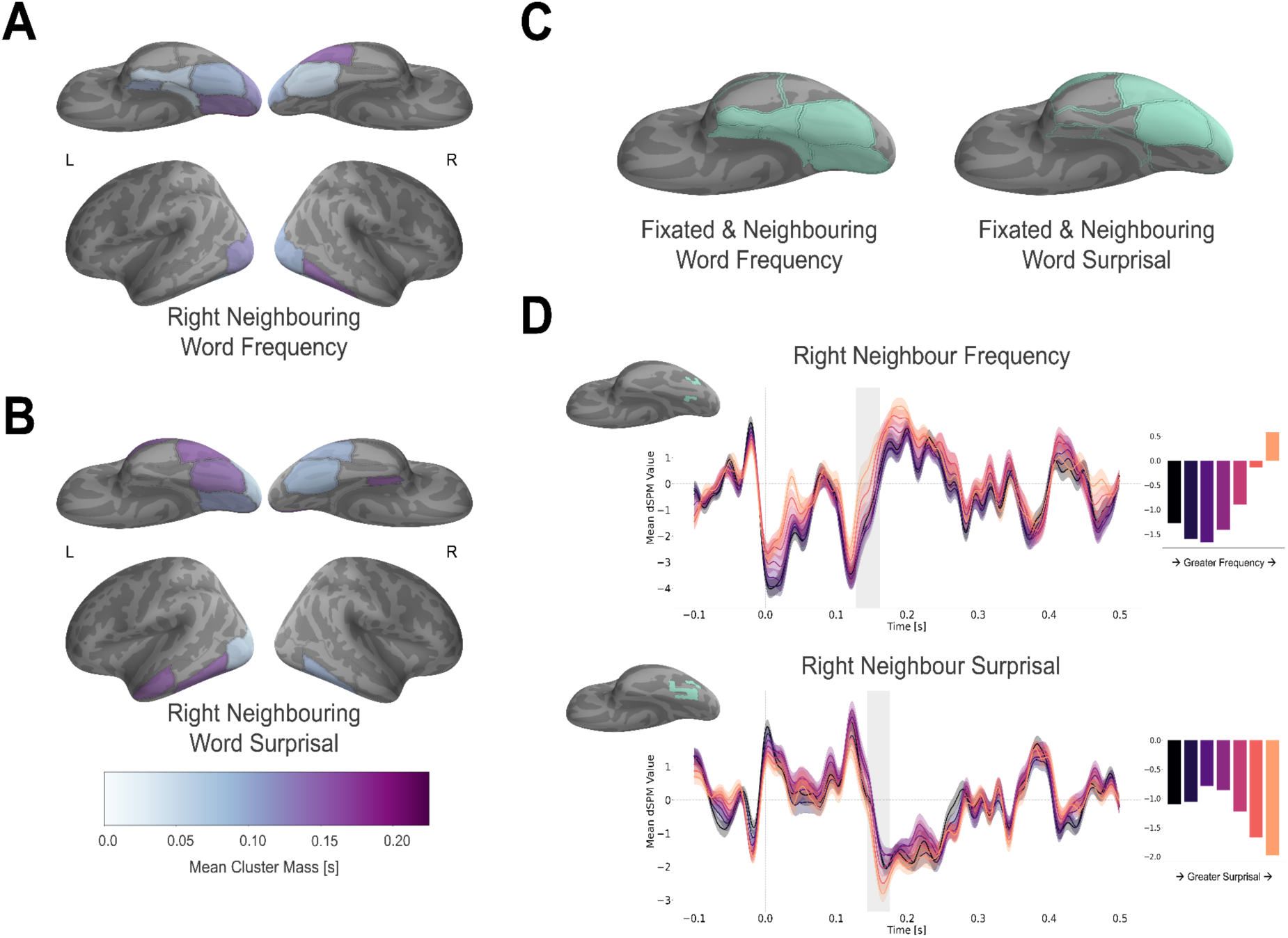
The influence of right neighbour properties. (A) Significant effects of the right neighbouring word’s frequency, 0-225 ms following the onset of first-pass first fixation on the foveal word. (B) Significant effects of the right neighbour’s surprisal during the same time window. (C) Regions that show overlapping effects of the fixated and neighbouring words’ frequency (left) and surprisal (right) in the 0-225 ms analysis window. (D) Time courses of activation for levels of right neighbour frequency (top; cluster: 128-161 ms) and surprisal (bottom; cluster: 144-175 ms) from significant clusters in the left posterior fusiform gyrus. The highlighted areas to the top-left of each time course display constituent sources within the ROI that were part of significant clusters overlapping with the shaded time window.

These findings are consistent with previous reports of parafoveal-on-foveal influences on neural activity, including parafoveal effects that emerge as early as 100 ms post-fixation onset (Degno et al., 2019; Pan et al., 2024). The overlapping spatial and temporal influences of both words’ properties provides novel evidence that the left hemisphere’s occipitotemporal visual word system (Fig. 4C) processes foveal and parafoveal words in parallel during natural reading. The strongest evidence for parallel processing was found in the left posterior fusiform gyrus (Figure 3, bottom-row, and Figure 4D), where activity during the first fixation on the foveal word was influenced by that word’s frequency and surprisal, as well as the right neighbouring word’s frequency and surprisal. These effects were found while accounting for word length, the amplitude of the incoming saccade, the length of the right neighbouring word, and the subsequent saccade in the deconvolution model. Although the left posterior fusiform clusters that showed neighbouring frequency and surprisal effects did not perfectly overlap in space, they did show near-simultaneous influences of the two properties in time (right frequency: 128-161 ms, right surprisal: 144-175 ms).

### 3.5 Neighbouring word frequency effects emerge independent of surprisal

The presence of the neighbouring word effects raised the question of whether the putatively independent influences of a word’s frequency and predictability begin while that word is still a parafoveal neighbour. Such a processing mechanism could explain the earliness of independent foveal word frequency and surprisal effects found in Section 3.2, shown in Figure 3 (beginning as early as 30 ms after fixation), as well as the effects of the two properties on eye movement behaviours that capture early stages of processing (i.e., first-pass first fixation durations and skipping probability).

To examine this further, we again split fixations into two categories, this time based on the 35th and 65th percentiles of surprisal of the *right neighbour*. We then tested for effects of the right neighbour’s frequency, in each category, during the 225 ms after onset of the fixation on the current word. Significant effects of frequency, in both surprisal analyses, which co-localized with effects of parafoveal surprisal, would provide additional evidence for local independent influences of the two properties, beginning before a word is fixated in foveal vision.

Four left hemisphere areas, and one right hemisphere area, showed main effects of right word surprisal, and effects of right word frequency independent of right word surprisal: the left lateral occipital cortex, lingual gyrus, posterior inferior temporal gyrus, and anterior middle temporal gyrus; and the right posterior fusiform gyrus. Of these areas, main effects of right word frequency, across all examined fixations, were found in the left lateral occipital cortex, left lingual gyrus, and right posterior fusiform gyrus (see Supplementary Table 6).

In the lateral occipital cortex, temporally overlapping effects of the right neighbour’s frequency emerged in clusters that began approximately 70 ms after the onset of the current fixation and showed obvious uniformity in both surprisal contexts (see Figure 5). In the left lingual gyrus, similarly timed effects of right neighbour frequency emerged closer to 200 ms after the onset of the fixation. Although these appeared to show reversed patterns with increasing frequency in high and low surprisal contexts, the clusters were not overlapping in space and the multiple sources of signal polarity in underlying dipolar generators of the MEG source estimates makes it difficult to assign functional interpretation to higher/lower estimated amplitudes (see Section 2.5.4). In the right posterior fusiform, localized responses were more variable and did not show putative visual response peaks (see Supplementary Figure 1), as were clear in the left hemisphere ROIs.

**Figure 5.**
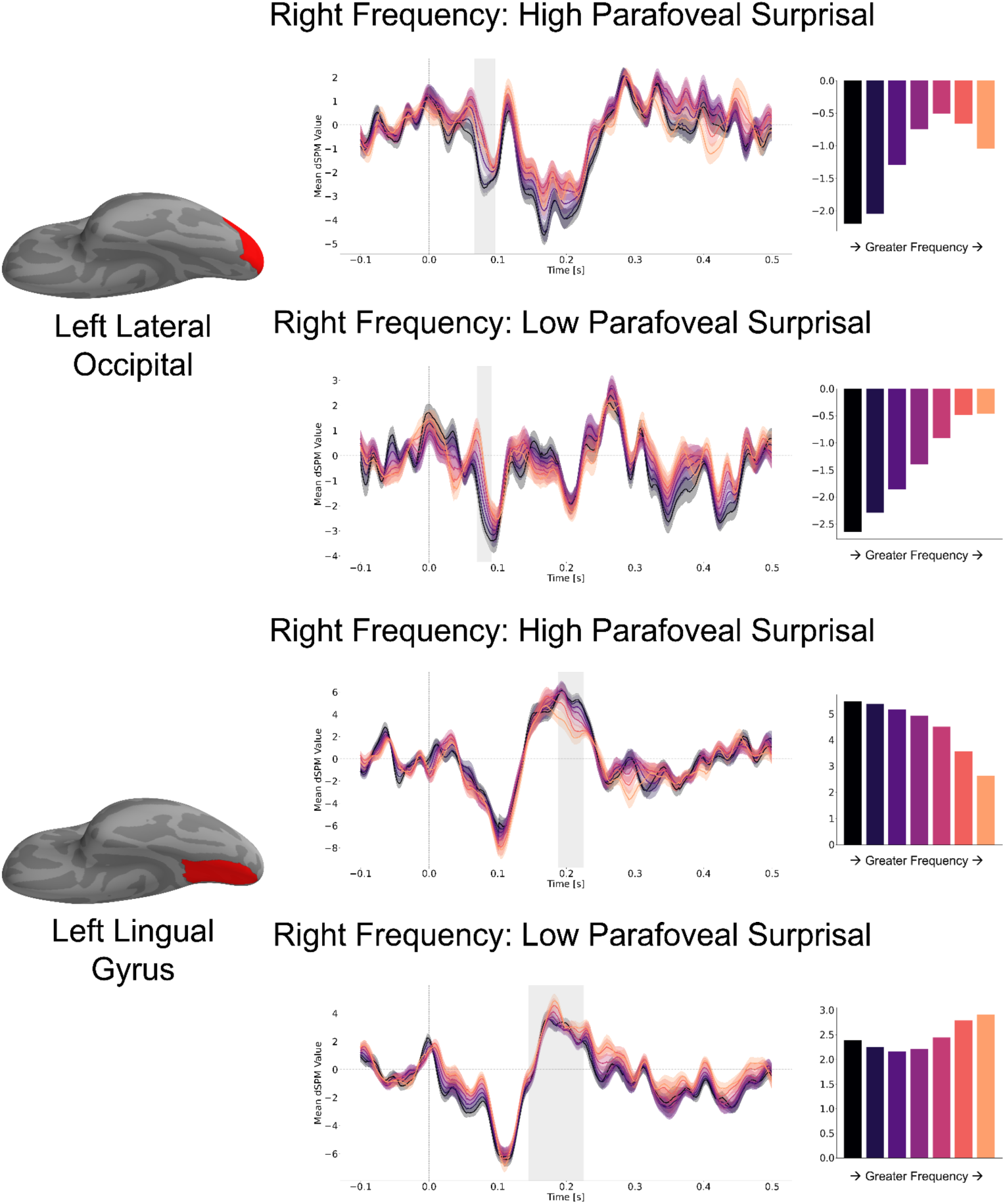
Significant influences of the right neighbouring word’s frequency, regardless of whether that word was of high/low surprisal, in the left lateral occipital cortex (high surprisal frequency effects: 66-96 ms; low surprisal frequency effects: 70-91 ms) and left lingual gyrus (high surprisal frequency effects: 188-225 ms; low surprisal frequency effects: 145-225 ms). The 0 milliseconds time point indicates the onset of the fixation on the preceding word. For each region, clusters that showed similarly timed frequency effects across high/low surprisal were selected for visualization. The visualized clusters within each ROI were not defined from the same set of constituent sources, which may account for the reversal of polarity with increasing frequency observed across high and low surprisal in the lingual gyrus.

We did not find right word frequency effects in both levels of surprisal within the left posterior fusiform ROI that showed near influences of the two properties across all examined fixations (Figure 4D). While this means this ROI did not meet our previously established criterion for the strongest evidence of local independent effects, the two ROIs that did meet this criterion were the posterior fusiform’s immediate neighbours: the left lateral occipital cortex and lingual gyrus.

We also did not find clear evidence for independent impacts of the right neighbouring word’s frequency and surprisal in those ROIs that showed these effects on the fixated word at very early time points (i.e., the left parahippocampal cortex and anterior inferior temporal gyrus), where effects emerged within 30 ms after fixation onset; see Figure 3. Instead, overlapping effects of the neighbouring word’s frequency and surprisal were predominantly found in more posterior areas, including the bilateral lateral occipital cortex and lingual gyrus, suggesting that information about a word’s frequency and surprisal may be relayed more anteriorly as it is brought into foveal vision.

In summary, several left occipitotemporal ROIs displayed parafoveal effects of the right neighbouring word’s frequency and surprisal. Amongst these, the left posterior fusiform demonstrated evidence for parallel visual processing, with common effects of four examined properties (current/neighbouring frequency and surprisal). The left lingual gyrus and left lateral occipital cortex both displayed effects of the right neighbouring word’s frequency, regardless of whether that word was of high or low surprisal, and main effects of the right neighbouring word’s surprisal, within 200 ms after the onset of the current fixation. This suggests that both properties began independently influencing neural activity in these ROIs prior to fixations on the word.

### 3.6 Parafoveal effects of frequency and surprisal distinguish subsequent word skipping

Having established the parafoveal effects of frequency and surprisal, we asked whether each property’s influence on neural activity differed when the relevant word was subsequently skipped or fixated. We repeated the analysis of neural activity from 0-225 ms after fixation onset, separately modelling the influences of all properties on neural activity elicited by first-pass first fixations that resulted in each case.

As shown in Figure 6 (cluster details in Supplementary Tables 7 & 8), significant effects of the right neighbouring words were more widespread when that word was subsequently skipped compared to when it was subsequently fixated. Parafoveal frequency effects that only emerged when the subsequent word was skipped were found in bilateral posterior fusiform and lingual gyri, the left lateral occipital cortex and posterior middle temporal gyrus, and the right anterior inferior temporal gyrus. Parafoveal surprisal effects that emerged exclusively when the subsequent word was skipped were found in the bilateral middle temporal gyrus (both anterior and posterior) and lateral occipital cortex, as well as the left posterior fusiform gyrus. The left lateral occipital cortex, left posterior fusiform, and left posterior middle temporal gyrus all showed both frequency and surprisal effects of the parafoveal word, that only emerged ahead of it being skipped. Post-hoc repeated measures ANOVAs on mean cluster activity, with the factors of skipped versus fixated and parafoveal frequency or parafoveal surprisal, revealed a significant interaction with parafoveal frequency in the left posterior fusiform (13-51 ms, F(6, 391) = 2.77, p = 0.014) and a significant interaction with parafoveal surprisal in the left middle temporal gyrus, both anteriorly (160-201 ms, F(6, 391) = 3.27, p = 0.005) and posteriorly (63-85 ms; F(6, 391) = 2.88, p = 0.011).

**Figure 6.**
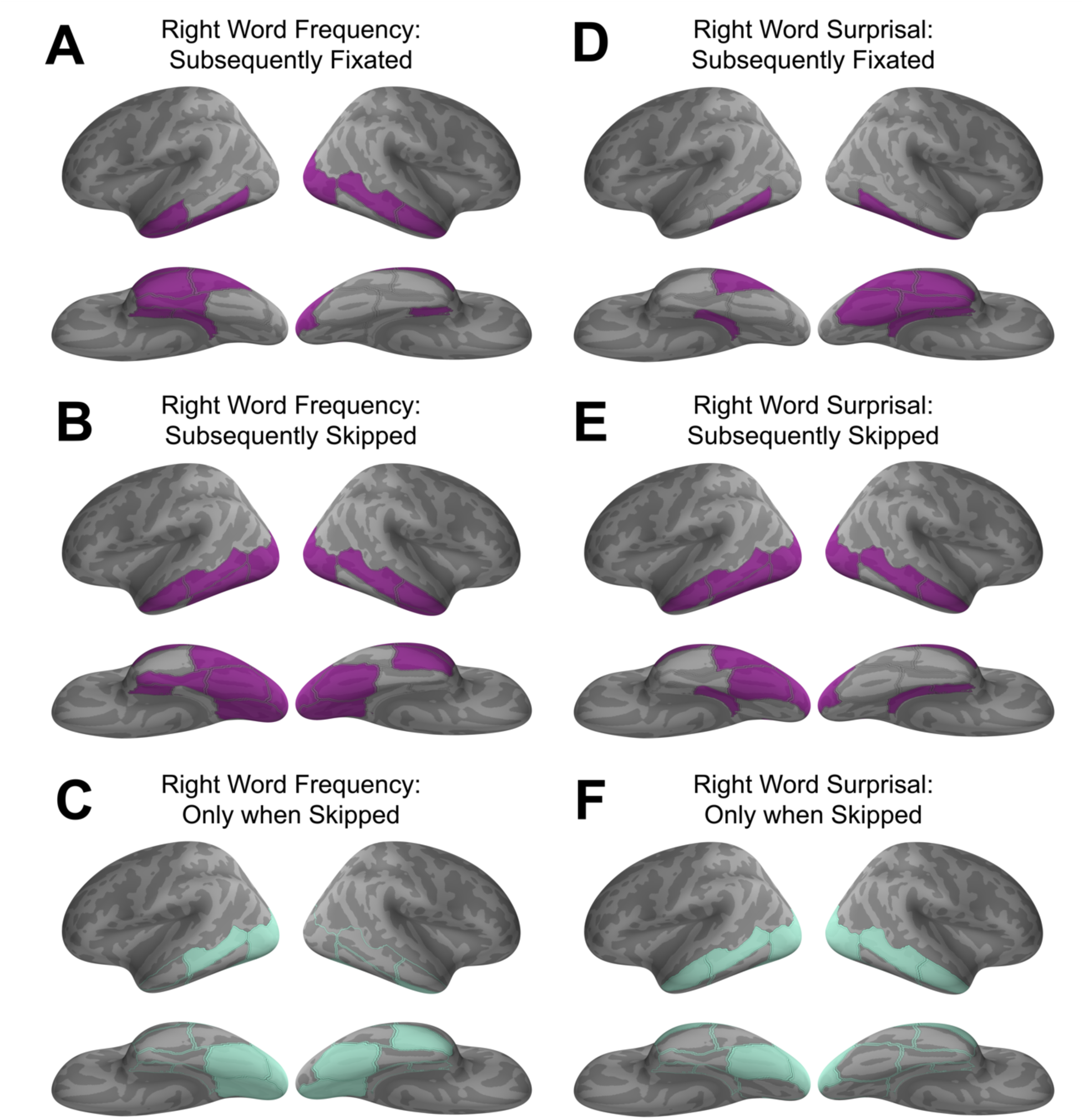
Effects of the right neighbouring word, on the current fixation, when that word was subsequently fixated or skipped. Left column: ROIs showing significant effects of right neighbour frequency when subsequently fixated (A), subsequently skipped (B), and those ROIs that only showed frequency effects when the word was subsequently skipped (C). Right column: ROIs showing significant effects of right neighbour surprisal when subsequently fixated (D), subsequently skipped (F), and those ROIs that only showed surprisal effects when the word was subsequently skipped (E).

### 3.7 Late interactions emerge in lateral and anterior temporal areas

Finally, as summarized in the Introduction, previous M/EEG studies found interactions between frequency and surprisal on the N400 ERP component (e.g., Kretzschmar et al., 2015; Payne et al., 2015; Alday et al., 2017), where frequency effects were diminished or absent when words were highly predicted by their preceding context. In a final set of exploratory analyses, we tested whether these previously reported interactions between frequency and surprisal emerged in later, deconvolved activity after the onset of fixations, corresponding to the typical N400 time window (225-500 ms post fixation onset).

As shown in Figure 7A, and in contrast to the 0-225 ms results shown in Figure 1, relatively few areas contained effects of both frequency and surprisal, across all examined fixations, in the 225-500 ms window. These areas were the bilateral lateral occipital cortex, and the right posterior inferior temporal gyrus, cuneus, and pericalcarine cortex. In contrast, 10 examined ROIs (Figure 7B) showed the multiplicative pattern found in previous studies, with significant effects of word frequency emerging in high surprise (low predictability) contexts but not those of low surprise (high predictability). These areas were predominantly found in anterior temporal or middle temporal areas, including the bilateral anterior fusiform and inferior temporal gyri, and the posterior middle temporal gyrus. Additional ROIs matching the pattern were the left posterior fusiform and parahippocampal cortex, and the right posterior inferior temporal gyrus and entorhinal cortex. Cluster information is reported in Supplementary Table 9.

**Figure 7.**
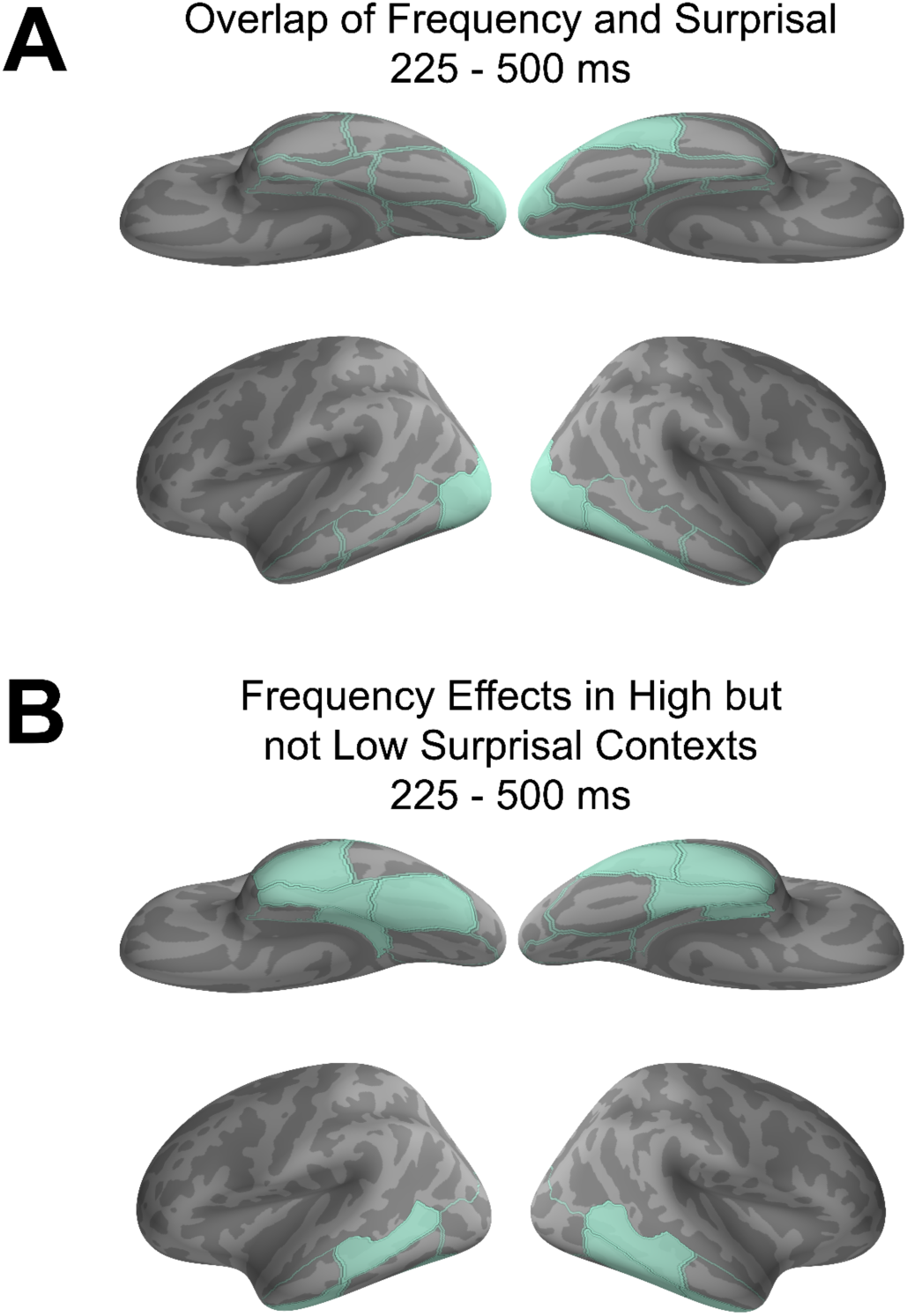
(A): Five ROIs displayed significant effects of both fixated word frequency and fixated word surprisal in the 225-500 ms post-fixation analysis window: the bilateral lateral occipital cortex, the right posterior inferior temporal gyrus, right cuneus, and right pericalcarine cortex. (B): Ten ROIs displayed significant effects of word frequency in only high surprisal (low predictive) contexts: the bilateral anterior fusiform and inferior temporal gyri, the bilateral posterior middle temporal gyrus, the left posterior fusiform gyrus, left parahippocampal cortex, the right posterior inferior temporal gyrus and the right entorhinal cortex.

We evaluated the multiplicative pattern further by conducting ANOVAs (frequency x surprisal category) on the mean activity within clusters in each of these ROIs. Six of the identified ROIs showed significant interactions between frequency and surprisal category. These were the left anterior fusiform gyrus (F(6,391) = 2.91, p = 0.01), right anterior inferior temporal gyrus (F(6, 391) = 2.16, p = 0.05), right posterior inferior temporal gyrus (F(6, 391) = 2.31, p = 0.036), right entorhinal cortex (F(6, 391) = 2.30, p = 0.037), and both the left (F(6, 391) = 4.54, p < 0.001) and right (F(6, 391) = 3.22, p = 0.005) posterior middle temporal gyri. Three ROIs, the left anterior inferior temporal gyrus, left parahippocampal cortex, and right anterior fusiform, showed main effects of word frequency in the ANOVA, but not significant interactions (all p-values > 0.150). The left posterior fusiform showed neither significant main effects nor a significant interaction (F(6, 391) = 1.69, p = 0.128), despite a significant effect of frequency in a one-way ANOVA conducted on the high surprisal responses (F(6,195) = 2.88, p = 0.011).

## 4. DISCUSSION

These results provide a spatiotemporal map of how the brain recognizes and integrates words during visual reading, and new insight into how linguistic properties of those words come to influence eye movements. In doing so, the results solve a long-standing discrepancy (see Kretzschmar et al., 2015; Sereno et al., 2020; Huizeling et al., 2022): although word frequency and predictability exert additive effects on fixation durations (e.g., Staub & Benatar, 2013; Staub, 2015), neural activity patterns that could explain those effects had remained unidentified.

Here, we reported what we believe to be the first localization of independent frequency and predictability effects during visual reading, which emerged within 225 ms of fixation onsets and matched the observed influences on eye movements. Our results provide several new details regarding these influences: (1) they appear in parafoveal processing, in left posterior visual and occipital areas during fixation on the preceding word; (2) they shift anteriorly within left ventral temporal cortex, when a word is fixated in foveal vision; and (3) across this posterior-to-anterior shift, the effects emerge early enough to influence fixation durations and saccade planning.

Beyond this, we replicated prior findings showing attenuation of frequency effects on later (225-500 ms) neural responses in constraining contexts. This is consistent with predictive coding accounts of the N400 (Wang et al., 2018, 2025; Nour Eddine et al., 2024) and adds further validity to the current results. We also provided new evidence suggesting that parafoveal words are processed to a greater degree, particularly in lateral and ventral temporal areas, prior to being skipped. Altogether, this grounds several decades of attested eye-movement patterns in localized brain activity. We discuss each of the primary findings in further detail below.

### 4.1 Word frequency and surprisal effects are grounded in left occipitotemporal activity

Matching the additive influences on fixation durations, several left ventral and occipitotemporal areas housed independent effects of frequency and predictability. The strongest evidence was found in the left fusiform, the inferior temporal (ITG) and parahippocampal gyri, as well as the right lateral occipital cortex. Here, fixated word frequency affected neural responses within 225 ms after fixation onset alongside separate effects of fixated word surprisal.

These findings match the results of several previous studies, though none had yet demarcated this pattern in space and time during natural reading. Prior M/EEG investigations and studies using intracranial recordings reported frequency and/or context effects as early as 120 milliseconds after word presentation, often in the same occipitotemporal areas observed here (Sereno et al., 1998; Sereno et al., 2003; Dambacher et al., 2006; Penolazzi et al., 2007; Gwilliams et al., 2016; Sereno et al., 2020; Woolnough et al, 2020; Flick et al., 2021). fMRI studies of natural reading have also found frequency (Schuster et al., 2016, 2025, although see Desai et al., 2020) and word context effects (Henderson et al., 2016; Schuster et al., 2016, 2020, 2021) in left occipitotemporal cortex.

One notable feature of our results is the early latency at which effects emerged in anterior temporal areas. These latencies (< 30 ms after fixation onset) are earlier than the time at which we expect visual information to activate primary cortex and be relayed anteriorly (Tarkiainen et al., 1998; Thorpe, Fize, & Marlot, 1998). This appears to be reconciled by analyses examining parafoveal processing of the subsequent word, showing that frequency and surprisal information were extracted prior to fixation, with independent effects emerging in posterior areas (i.e., the left lateral occipital cortex and lingual gyrus). In the lateral occipital cortex, parafoveal effects emerged on the first apparent peak of responses, at approximately 100 ms post-fixation (see Figure 5). This is consistent with a recent co-registered EEG and eye-tracking study, which showed that parafoveal information was extracted within 100 ms and facilitated very early (< 70 ms) foveal processing of the next word (Degno et al., 2019). Recent combined MEG and eye-tracking research has also evidenced parafoveal word recognition and integration, using factorial manipulations of select target words in sentence contexts (Pan et al., 2023, 2024).

These patterns suggest that information about a parafoveal word’s frequency and surprisal is simultaneously taken in by early visual and occipitotemporal areas (lateral occipital cortex, lingual gyrus, posterior fusiform gyrus; Figure 8) upon the start of a fixation. This processing is then shifted more anteriorly while another bottom-up pass of information about that word is received from foveal vision, once fixated, alongside information about the *next* parafoveal word. Our findings further suggest that the left posterior fusiform plays a central role in integrating the sources and maintenance of parafoveal and foveal information, as this area was the only region to show effects of all four properties of the fixated and neighbouring words.

**Figure 8.**
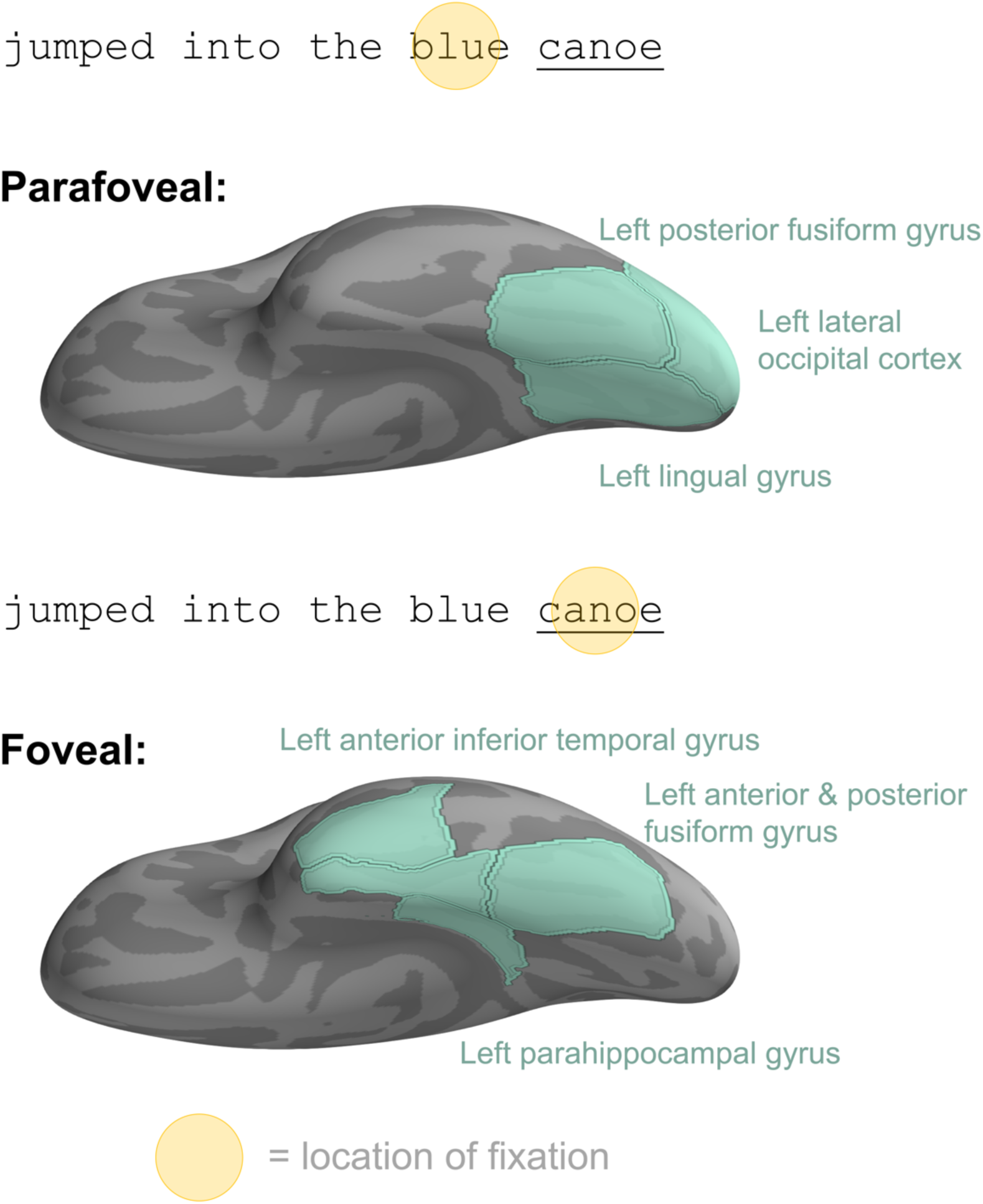
Summary of left ventral and occipitotemporal areas that housed locally independent effects of word frequency and surprisal parafoveally (top) and once fixated (bottom), matching the influences of these properties on first fixation durations. The example highlights regions related to the word canoe when fixating on the preceding word (blue; top) or the word itself (bottom). The left posterior fusiform gyrus showed effects of both properties in both parafoveal and foveal processing. It also appeared to distinguish between parafoveal words that are subsequently skipped or fixated, based on effects of word frequency.

### 4.2 Parafoveal frequency and surprisal effects precede word skipping

While length is known to be a predictor of whether or not a word will be skipped (Rayner & McConkie, 1976; Brysbaert & Vitu, 1998), eye-tracking research has demonstrated that both word frequency (Rayner, Sereno, & Raney, 1996) and predictability (Ehrlich & Rayner, 1981; Balota, Pollatsek, & Rayner, 1985; Rayner & Well, 1996; Drieghe, Rayner, & Pollatsek, 2005) influence skipping probabilities when target words are length-matched. Our results link these effects to parafoveal influences on neural activity. Overall, the current finding of widespread parafoveal effects before skips is consistent with a previous fMRI study of natural reading, which found greater activation ahead of skipped words in bilateral occipital cortex and the left posterior middle temporal gyrus (Schuster et al., 2016). The same study reported effects of word frequency in the left occipitotemporal cortex and predictability in middle and superior temporal gyri. This closely matches the skipping results found here, with our improved temporal resolution demonstrating that these patterns emerge within 225 ms of the preceding fixation onset.

Our results specifically highlight the left posterior fusiform and left middle temporal gyrus as playing important roles in visual word processing that facilitates readers’ abilities to skip over approximately one-third of all words (Rayner, 1998). Previous studies have implicated the left fusiform and nearby cortex in the look-up of a visual word’s meaning and pronunciation (i.e., its entry in one’s mental lexicon; Taylor et al., 2019; Woolnough et al., 2020), while left lateral temporal responses have been implicated in both lexical access and compositional processing (Fruchter et al., 2015; Flick & Pylkkänen, 2020). Although more work is needed to precisely identify the functions of each of these areas in word skipping, the past results suggest that the left fusiform may perform an initial look-up of the parafoveal word’s meaning, while the left middle temporal gyrus assesses the contextual fit of that word with the prior context; both of which influence the decision to skip. These computations may be able to be implemented rapidly, enabling the skips, when words are short, high frequency, or highly predictable. If a parafoveal word is not skipped, the left posterior fusiform integrates foveal information from the fixation on that word, with the parafoveal information that was previously extracted. This is demonstrated by the significant effects of both parafoveal and foveal word properties in this region.

### 4.3 Limitations & Future Directions

There are limitations to the current work, which we hope to refine in future research. First, the spatial resolution of MEG precludes us from drawing strong conclusions about the precise localization of frequency and surprisal effects. For instance, past results have shown that responses localized to anterior temporal areas are not necessarily only generated in that region (Hauk et al., 2011). Second, we did not comprehensively analyze regressions, re-fixations, or fixations on closed class words, all of which must be considered in a complete neuroscientific account of reading. Finally, there remain important questions about the causality of correlational effects observed here, particularly related to the influence of neural activity on eye movements. The pursuit of these answers would likely benefit from future work using stimulation methods and/or intracranial recordings during natural reading.

## 5. Conclusion

In this work, we have attempted to provide a spatiotemporal characterization of how visual reading unfolds on a fixation-by-fixation basis. The findings suggest that information about an upcoming word’s meaning and fit with the preceding context is extracted by a set of early visual and occipitotemporal areas – including the left lingual gyrus, posterior fusiform gyrus, and lateral occipital cortex – upon fixation on the preceding word. By the time the eyes fixate the next word in foveal vision, this information is encoded in more anterior ventral areas and continues to be extracted by the left posterior fusiform, in parallel with information about the *next* parafoveal word. The timing of the effects suggests that this occipitotemporal system is the first port of call where neural activity exerts downstream influences on fixation durations. We also show that patterns of neural activity in the left fusiform and middle temporal gyri distinguish whether an upcoming word will be skipped or fixated. These results precisely map out the dynamics of neural activity in left occipitotemporal cortex that recognize and integrate visual words in natural reading.

## Acknowledgements

We thank Jacqueline Fallon for assistance with data collection. We thank Brenden Lake, Alec Marantz, David Poeppel, and Alex White for comments on earlier drafts of this manuscript. We also thank the organizers and audiences of the 2022 Leading Edge Workshop on Co-registration. This work was supported by a Henry M. MacCracken Doctoral Fellowship from New York University to G.F. and a Natural Sciences and Engineering Research Council of Canada Postdoctoral Fellowship to G.F. It was also supported by the National Science Foundation (Award #2335767) to L.P. and Award G1001 from the NYUAD Institute, New York University Abu Dhabi to L.P.

## References

Adachi, Y., Shimogawara, M., Higuchi, M., Haruta, Y., & Ochiai, M. (2001). Reduction of non-periodic environmental magnetic noise in MEG measurement by continuously adjusted least squares method. IEEE Transactions on Applied Superconductivity, 11(1), 669–672. 10.1109/77.919433

Alday, P. M., Schlesewsky, M., & Bornkessel-Schlesewsky, I. (2017). Electrophysiology Reveals the Neural Dynamics of Naturalistic Auditory Language Processing: Event-Related Potentials Reflect Continuous Model Updates. Eneuro, 4(6), ENEURO.0311-16.2017. 10.1523/ENEURO.0311-16.2017

Baillet, S. (2017). Magnetoencephalography for brain electrophysiology and imaging. Nature Neuroscience, 20(3), 327–339. 10.1038/nn.4504

Balota, D. A., Pollatsek, A., & Rayner, K. (1985). The interaction of contextual constraints and parafoveal visual information in reading. Cognitive Psychology, 17(3), 364–390. 10.1016/0010-0285(85)90013-1

Balota, D. A., Yap, M. J., Hutchison, K. A., Cortese, M. J., Kessler, B., Loftis, B., Neely, J. H., Nelson, D. L., Simpson, G. B., & Treiman, R. (2007). The English Lexicon Project. Behavior Research Methods, 39(3), 445–459. 10.3758/BF03193014

Bates, D., Mächler, M., Bolker, B., & Walker, S. (2015). Fitting Linear Mixed-Effects Models Using lme4. Journal of Statistical Software, 67, 1–48. 10.18637/jss.v067.i01

Benjamini, Y., & Yekutieli, D. (2001). The Control of the False Discovery Rate in Multiple Testing under Dependency. The Annals of Statistics, 29(4), 1165–1188.

Brainard, D. H. (1997). The Psychophysics Toolbox. Spatial Vision, 10(4), 433–436. 10.1163/156856897X00357

Brysbaert, M., & Vitu, F. (1998). Chapter 6 - Word Skipping: Implications for Theories of Eye Movement Control in Reading. In G. Underwood (Ed.), Eye Guidance in Reading and Scene Perception (pp. 125–147). Elsevier Science Ltd. 10.1016/B978-008043361-5/50007-9

Caliskan, N., Milligan, S., & Schotter, E. R. (2023). Readers scrutinize lexical familiarity only in the absence of expectations: Evidence from lexicality effects on event-related potentials. Brain and Language, 238, 105232. 10.1016/j.bandl.2023.105232

Caucheteux, C., Gramfort, A., & King, J.-R. (2022). Deep language algorithms predict semantic comprehension from brain activity. Scientific Reports, 12(1), 16327. 10.1038/s41598-022-20460-9

Choi, W., Desai, R. H., & Henderson, J. M. (2014). The neural substrates of natural reading: a comparison of normal and nonword text using eyetracking and fMRI. Frontiers in human neuroscience, 8, 1024. 10.3389/fnhum.2014.01024

Coco, M. I., Nuthmann, A., & Dimigen, O. (2020). Fixation-related brain potentials during semantic integration of object–scene information. Journal of Cognitive Neuroscience, 32(4), 571–589.

Dale, A. M., Fischl, B., & Sereno, M. I. (1999). Cortical Surface-Based Analysis: I. Segmentation and Surface Reconstruction. NeuroImage, 9(2), 179–194. 10.1006/nimg.1998.0395

Dale, A. M., Liu, A. K., Fischl, B. R., Buckner, R. L., Belliveau, J. W., Lewine, J. D., & Halgren, E. (2000). Dynamic Statistical Parametric Mapping: Combining fMRI and MEG for High-Resolution Imaging of Cortical Activity. Neuron, 26(1), 55–67. 10.1016/S0896-6273(00)81138-1

Dambacher, M., Kliegl, R., Hofmann, M., & Jacobs, A. M. (2006). Frequency and predictability effects on event-related potentials during reading. Brain Research, 1084(1), 89–103. 10.1016/j.brainres.2006.02.010

Degno, F., Loberg, O., Zang, C., Zhang, M., Donnelly, N., & Liversedge, S. P. (2019). Parafoveal previews and lexical frequency in natural reading: Evidence from eye movements and fixation-related potentials. Journal of experimental psychology. General, 148(3), 453–474. 10.1037/xge0000494

Degno, F., & Liversedge, S. P. (2020). Eye Movements and Fixation-Related Potentials in Reading: A Review. Vision, 4(1), Article 1. 10.3390/vision4010011

Dehaene, S., Pegado, F., Braga, L. W., Ventura, P., Filho, G. N., Jobert, A., Dehaene-Lambertz, G., Kolinsky, R., Morais, J., & Cohen, L. (2010). How Learning to Read Changes the Cortical Networks for Vision and Language. Science, 330(6009), 1359–1364. 10.1126/science.1194140

Desai, R. H., Choi, W., & Henderson, J. M. (2020). Word frequency effects in naturalistic reading. Language, Cognition and Neuroscience, 35(5), 583–594. 10.1080/23273798.2018.1527376

Desikan, R. S., Ségonne, F., Fischl, B., Quinn, B. T., Dickerson, B. C., Blacker, D., Buckner, R. L., Dale, A. M., Maguire, R. P., Hyman, B. T., Albert, M. S., & Killiany, R. J. (2006). An automated labeling system for subdividing the human cerebral cortex on MRI scans into gyral based regions of interest. NeuroImage, 31(3), 968–980. 10.1016/j.neuroimage.2006.01.021

Dikker, S., Rabagliati, H., & Pylkkänen, L. (2009). Sensitivity to syntax in visual cortex. Cognition, 110(3), 293–321. 10.1016/j.cognition.2008.09.008

Dimigen, O., Sommer, W., Hohlfeld, A., Jacobs, A. M., & Kliegl, R. (2011). Coregistration of eye movements and EEG in natural reading: Analyses and review. Journal of Experimental Psychology: General, 140(4), 552–572. 10.1037/a0023885

Drieghe, D., Rayner, K., & Pollatsek, A. (2005). Eye Movements and Word Skipping During Reading Revisited. Journal of Experimental Psychology: Human Perception and Performance, 31(5), 954–969. 10.1037/0096-1523.31.5.954

Eddine, S. N., Brothers, T., Wang, L., Spratling, M., & Kuperberg, G. R. (2024). A predictive coding model of the N400. Cognition, 246, 105755. 10.1016/j.cognition.2024.105755

Ehinger, B. V., & Dimigen, O. (2019). Unfold: An integrated toolbox for overlap correction, non-linear modeling, and regression-based EEG analysis. PeerJ, 7, e7838. 10.7717/peerj.7838

Ehrlich, S. F., & Rayner, K. (1981). Contextual effects on word perception and eye movements during reading. Journal of Verbal Learning and Verbal Behavior, 20(6), 641–655. 10.1016/S0022-5371(81)90220-6

Engemann, D. A., & Gramfort, A. (2015). Automated model selection in covariance estimation and spatial whitening of MEG and EEG signals. NeuroImage, 108, 328–342. 10.1016/j.neuroimage.2014.12.040

Fischl, B., Sereno, M. I., Tootell, R. B. H., & Dale, A. M. (1999). High-resolution intersubject averaging and a coordinate system for the cortical surface. Human Brain Mapping, 8(4), 272–284. 10.1002/(SICI)1097-0193(1999)8:4<272::AID-HBM10>3.0.CO;2-4

Fischl, B., van der Kouwe, A., Destrieux, C., Halgren, E., Ségonne, F., Salat, D. H., Busa, E., Seidman, L. J., Goldstein, J., Kennedy, D., Caviness, V., Makris, N., Rosen, B., & Dale, A. M. (2004). Automatically Parcellating the Human Cerebral Cortex. Cerebral Cortex, 14(1), 11–22. 10.1093/cercor/bhg087

Flick, G., Abdullah, O., & Pylkkänen, L. (2021). From letters to composed concepts: A magnetoencephalography study of reading. Human Brain Mapping, 42(15), 5130–5153. 10.1002/hbm.25608

Flick, G., & Pylkkänen, L. (2020). Isolating syntax in natural language: MEG evidence for an early contribution of left posterior temporal cortex. Cortex, 127, 42–57. 10.1016/j.cortex.2020.01.025

Fruchter, J., Linzen, T., Westerlund, M., & Marantz, A. (2015). Lexical Preactivation in Basic Linguistic Phrases. Journal of Cognitive Neuroscience, 27(10), 1912–1935. 10.1162/jocn_a_00822

Goldstein, A., Grinstein-Dabush, A., Schain, M., Wang, H., Hong, Z., Aubrey, B., Nastase, S. A., Zada, Z., Ham, E., Feder, A., Gazula, H., Buchnik, E., Doyle, W., Devore, S., Dugan, P., Reichart, R., Friedman, D., Brenner, M., Hassidim, A., … Hasson, U. (2024). Alignment of brain embeddings and artificial contextual embeddings in natural language points to common geometric patterns. Nature Communications, 15(1), 2768. 10.1038/s41467-024-46631-y

Goldstein, A., Zada, Z., Buchnik, E., Schain, M., Price, A., Aubrey, B., Nastase, S. A., Feder, A., Emanuel, D., Cohen, A., Jansen, A., Gazula, H., Choe, G., Rao, A., Kim, C., Casto, C., Fanda, L., Doyle, W., Friedman, D., … Hasson, U. (2022). Shared computational principles for language processing in humans and deep language models. Nature Neuroscience, 25(3), 369–380. 10.1038/s41593-022-01026-4

Grainger, J., Dufau, S., & Ziegler, J. C. (2016). A Vision of Reading. Trends in Cognitive Sciences, 20(3), 171–179. 10.1016/j.tics.2015.12.008

Gramfort, A., Luessi, M., Larson, E., Engemann, D. A., Strohmeier, D., Brodbeck, C., Goj, R., Jas, M., Brooks, T., Parkkonen, L., & Hämäläinen, M. (2013). MEG and EEG data analysis with MNE-Python. Frontiers in Neuroscience, 7. 10.3389/fnins.2013.00267

Gramfort, A., Luessi, M., Larson, E., Engemann, D. A., Strohmeier, D., Brodbeck, C., Parkkonen, L., & Hämäläinen, M. S. (2014). MNE software for processing MEG and EEG data. NeuroImage, 86, 446–460. 10.1016/j.neuroimage.2013.10.027

Gwilliams, L., Lewis, G. A., & Marantz, A. (2016). Functional characterisation of letter-specific responses in time, space and current polarity using magnetoencephalography. NeuroImage, 132, 320–333. 10.1016/j.neuroimage.2016.02.057

Hari, R., & Puce, A. (2023). MEG-EEG Primer. Oxford University Press.

Hauk, O., Wakeman, D. G., & Henson, R. (2011). Comparison of noise-normalized minimum norm estimates for MEG analysis using multiple resolution metrics. NeuroImage, 54(3), 1966–1974. 10.1016/j.neuroimage.2010.09.053

Henderson, J. M., Choi, W., Lowder, M. W., & Ferreira, F. (2016). Language structure in the brain: A fixation-related fMRI study of syntactic surprisal in reading. NeuroImage, 132, 293–300. 10.1016/j.neuroimage.2016.02.050

Henderson, J. M., Choi, W., Luke, S. G., & Desai, R. H. (2015). Neural correlates of fixation duration in natural reading: Evidence from fixation-related fMRI. NeuroImage, 119, 390–397. 10.1016/j.neuroimage.2015.06.072

Himmelstoss, N. A., Schuster,Sarah, Hutzler,Florian, Moran,Rosalyn, & and Hawelka, S. (2020). Co-registration of eye movements and neuroimaging for studying contextual predictions in natural reading. Language, Cognition and Neuroscience, 35(5), 595–612. 10.1080/23273798.2019.1616102

Huizeling, E., Arana, S., Hagoort, P., & Schoffelen, J.-M. (2022). Lexical Frequency and Sentence Context Influence the Brain’s Response to Single Words. Neurobiology of Language, 3(1), 149–179. 10.1162/nol_a_00054

Kennedy, A., Pynte, J., Murray, W. S., & Paul, S.-A. (2013). Frequency and predictability effects in the Dundee Corpus: An eye movement analysis. Quarterly Journal of Experimental Psychology, 66(3), 601–618. 10.1080/17470218.2012.676054

Kliegl, R., Grabner,Ellen, Rolfs,Martin, & and Engbert, R. (2004). Length, frequency, and predictability effects of words on eye movements in reading. European Journal of Cognitive Psychology, 16(1–2), 262–284. 10.1080/09541440340000213

Kragel, J. E., VanHaerents, S., Templer, J. W., Schuele, S., Rosenow, J. M., Nilakantan, A. S., & Bridge, D. J. (2020). Hippocampal theta coordinates memory processing during visual exploration. eLife, 9, e52108. 10.7554/eLife.52108

Kretzschmar, F., Schlesewsky, M., & Staub, A. (2015). Dissociating word frequency and predictability effects in reading: Evidence from coregistration of eye movements and EEG. *Journal of Experimental Psychology: Learning*, Memory, and Cognition, 41(6), 1648–1662. 10.1037/xlm0000128

Kutas, M., & Hillyard, S. A. (1980). Reading Senseless Sentences: Brain Potentials Reflect Semantic Incongruity. Science, 207(4427), 203–205. 10.1126/science.7350657

Kutas, M., & Hillyard, S. A. (1984). Brain potentials during reading reflect word expectancy and semantic association. Nature, 307(5947), 161–163. 10.1038/307161a0

Laszlo, S., & Federmeier, K. D. (2009). A beautiful day in the neighborhood: An event-related potential study of lexical relationships and prediction in context. Journal of Memory and Language, 61(3), 326–338. 10.1016/j.jml.2009.06.004

Lin, F.-H., Witzel, T., Ahlfors, S. P., Stufflebeam, S. M., Belliveau, J. W., & Hämäläinen, M. S. (2006). Assessing and improving the spatial accuracy in MEG source localization by depth-weighted minimum-norm estimates. NeuroImage, 31(1), 160–171. 10.1016/j.neuroimage.2005.11.054

Maris, E., & Oostenveld, R. (2007). Nonparametric statistical testing of EEG- and MEG-data. Journal of Neuroscience Methods, 164(1), 177–190. 10.1016/j.jneumeth.2007.03.024

Misra, K. (2022). *minicons: Enabling Flexible Behavioral and Representational Analyses of Transformer Language Models* (arXiv:2203.13112). arXiv. 10.48550/arXiv.2203.13112

Nikolaev, A. R., Meghanathan, R. N., & van Leeuwen, C. (2016). Combining EEG and eye movement recording in free viewing: Pitfalls and possibilities. Brain and Cognition, 107, 55–83. 10.1016/j.bandc.2016.06.004

Pan, Y., Frisson, S., Federmeier, K. D., & Jensen, O. (2024). Early parafoveal semantic integration in natural reading. eLife, 12, RP91327. 10.7554/eLife.91327

Pan, Y., Popov, T., Frisson, S., & Jensen, O. (2023). Saccades are locked to the phase of alpha oscillations during natural reading. PLOS Biology, 21(1), e3001968. 10.1371/journal.pbio.3001968

Payne, B. R., Lee, C., & Federmeier, K. D. (2015). Revisiting the incremental effects of context on word processing: Evidence from single-word event-related brain potentials. Psychophysiology, 52(11), 1456–1469. 10.1111/psyp.12515

Penolazzi, B., Hauk, O., & Pulvermüller, F. (2007). Early semantic context integration and lexical access as revealed by event-related brain potentials. Biological Psychology, 74(3), 374–388. 10.1016/j.biopsycho.2006.09.008

Radford, A., Wu, J., Child, R., Luan, D., Amodei, D., & Sutskever, I. (2019). Language models are unsupervised multitask learners. OpenAI blog, 1(8), 9.

Rayner, K. (1978). Eye movements in reading and information processing. Psychological Bulletin, 85(3), 618–660. 10.1037/0033-2909.85.3.618

Rayner, K. (1998). Eye movements in reading and information processing: 20 years of research. Psychological Bulletin, 124(3), 372–422. 10.1037/0033-2909.124.3.372

Rayner, K., Ashby, J., Pollatsek, A., & Reichle, E. D. (2004). The Effects of Frequency and Predictability on Eye Fixations in Reading: Implications for the E-Z Reader Model. Journal of Experimental Psychology: Human Perception and Performance, 30(4), 720–732. 10.1037/0096-1523.30.4.720

Rayner, K., & McConkie, G. W. (1976). What guides a reader’s eye movements? Vision Research, 16(8), 829–837. 10.1016/0042-6989(76)90143-7

Rayner, K., Sereno, S. C., & Raney, G. E. (1996). Eye movement control in reading: a comparison of two types of models. Journal of Experimental Psychology: Human Perception and Performance, 22(5), 1188

Rayner, K., & Well, A. D. (1996). Effects of contextual constraint on eye movements in reading: A further examination. Psychonomic Bulletin & Review, 3(4), 504–509. 10.3758/BF03214555

Schuster, S., Hawelka, S., Hutzler, F., Kronbichler, M., & Richlan, F. (2016). Words in Context: The Effects of Length, Frequency, and Predictability on Brain Responses During Natural Reading. Cerebral Cortex, 26(10), 3889–3904. 10.1093/cercor/bhw184

Schuster, S., Himmelstoss, N. A., Hutzler, F., Richlan, F., Kronbichler, M., & Hawelka, S. (2021). Cloze enough? Hemodynamic effects of predictive processing during natural reading. NeuroImage, 228, 117687. 10.1016/j.neuroimage.2020.117687

Schuster, S., Weiss, K.-L., Hutzler, F., Kronbichler, M., & Hawelka, S. (2025). Interactive and additive effects of word frequency and predictability: A fixation-related fMRI study. Brain and Language, 260, 105508. 10.1016/j.bandl.2024.105508

Sereno, S. C., Brewer, C. C., & O’Donnell, P. J. (2003). Context Effects in Word Recognition: Evidence for Early Interactive Processing. Psychological Science, 14(4), 328–333. 10.1111/1467-9280.14471

Sereno, S. C., Hand,Christopher J., Shahid,Aisha, Mackenzie,Ian G., & and Leuthold, H. (2020). Early EEG correlates of word frequency and contextual predictability in reading. Language, Cognition and Neuroscience, 35(5), 625–640. 10.1080/23273798.2019.1580753

Sereno, S. C., Rayner, K., & Posner, M. I. (1998). Establishing a time-line of word recognition: Evidence from eye movements and event-related potentials. NeuroReport, 9(10), 2195.

Smith, S. M., & Nichols, T. E. (2009). Threshold-free cluster enhancement: Addressing problems of smoothing, threshold dependence and localisation in cluster inference. NeuroImage, 44(1), 83–98. 10.1016/j.neuroimage.2008.03.061

Snell, J., & Grainger, J. (2019). Readers Are Parallel Processors. Trends in Cognitive Sciences, 23(7), 537–546. 10.1016/j.tics.2019.04.006

Staub, A. (2015). The Effect of Lexical Predictability on Eye Movements in Reading: Critical Review and Theoretical Interpretation. Language and Linguistics Compass, 9(8), 311–327. 10.1111/lnc3.12151

Staub, A., & Benatar, A. (2013). Individual differences in fixation duration distributions in reading. Psychonomic Bulletin & Review, 20(6), 1304–1311. 10.3758/s13423-013-0444-x

Tarkiainen, A., Helenius, P., Hansen, P. C., Cornelissen, P. L., & Salmelin, R. (1999). Dynamics of letter string perception in the human occipitotemporal cortex. Brain, 122(11), 2119–2132. 10.1093/brain/122.11.2119

Taylor, J. S. H., Davis, M. H., & Rastle, K. (2019). Mapping visual symbols onto spoken language along the ventral visual stream. Proceedings of the National Academy of Sciences, 116(36), 17723–17728. 10.1073/pnas.1818575116

Van Petten, C., & Kutas, M. (1990). Interactions between sentence context and word frequencyinevent-related brainpotentials. Memory & Cognition, 18(4), 380–393. 10.3758/BF03197127

Van Petten, C., & Kutas, M. (1991). Influences of semantic and syntactic context on open- and closed-class words. Memory & Cognition, 19(1), 95–112. 10.3758/BF03198500

Vinckier, F., Dehaene, S., Jobert, A., Dubus, J. P., Sigman, M., & Cohen, L. (2007). Hierarchical Coding of Letter Strings in the Ventral Stream: Dissecting the Inner Organization of the Visual Word-Form System. Neuron, 55(1), 143–156. 10.1016/j.neuron.2007.05.031

Wang, L., Kuperberg, G., & Jensen, O. (2018). Specific lexico-semantic predictions are associated with unique spatial and temporal patterns of neural activity. eLife, 7, e39061. 10.7554/eLife.39061

Wang, L., Nour Eddine, S., Brothers, T., Jensen, O., & Kuperberg, G. R. (2025). An implemented predictive coding model of lexico-semantic processing explains the dynamics of univariate and multivariate activity within the left ventromedial temporal lobe during reading comprehension. NeuroImage, 308, 120977. 10.1016/j.neuroimage.2024.120977

Winter, B. (2013). *Linear models and linear mixed effects models in R with linguistic applications* (arXiv:1308.5499). arXiv. 10.48550/arXiv.1308.5499

Wolf, T., Debut, L., Sanh, V., Chaumond, J., Delangue, C., Moi, A., Cistac, P., Rault, T., Louf, R., Funtowicz, M., Davison, J., Shleifer, S., von Platen, P., Ma, C., Jernite, Y., Plu, J., Xu, C., Le Scao, T., Gugger, S., … Rush, A. (2020). Transformers: State-of-the-Art Natural Language Processing. In Q. Liu & D. Schlangen (Eds.), Proceedings of the 2020 Conference on Empirical Methods in Natural Language Processing: System Demonstrations (pp. 38–45). Association for Computational Linguistics. 10.18653/v1/2020.emnlp-demos.6

Wood, S. N. (2017). Generalized Additive Models: An Introduction with R, Second Edition (2nd ed.). Chapman and Hall/CRC. 10.1201/9781315370279

Woolnough, O., Donos, C., Rollo, P. S., Forseth, K. J., Lakretz, Y., Crone, N. E., Fischer-Baum, S., Dehaene, S., & Tandon, N. (2021). Spatiotemporal dynamics of orthographic and lexical processing in the ventral visual pathway. Nature Human Behaviour, 5(3), 389–398. 10.1038/s41562-020-00982-w

