## Supplementary for "Reading ahead: Localized neural signatures of parafoveal word processing and skipping decisions"

Supplementary Table 1. Fixated Word Frequency Cluster Information, 0-225 ms

| Region | Max Cluster Duration [ms] | Max Cluster Sources | Corrected p-value |
|---|---|---|---|
| L aFus | 25 | 8 | 0.019 |
| L aITG | 21 | 7 | 0.020 |
| L pITG | 17 | 7 | 0.019 |
| L aMTG | 19 | 4 | 0.019 |
| L pMTG | 16 | 8 | 0.019 |
| L Lingual | 40 | 36 | 0.019 |
| L Pericalcarine | 37 | 9 | 0.019 |
| L Parahippocampal | 19 | 9 | 0.019 |
| L Entorhinal | 28 | 4 | 0.019 |
| R pFus | 23 | 13 | 0.019 |
| R aMTG | 12 | 5 | 0.024 |
| R pMTG | 21 | 3 | 0.039 |
| R Lateral Occip | 38 | 12 | 0.019 |
| R Pericalcarine | 23 | 5 | 0.019 |
| R Parahippocampal | 38 | 14 | 0.020 |
| R Entorhinal | 34 | 6 | 0.022 |

Abbreviations: a = anterior; p = posterior; Fus = fusiform gyrus; ITG = inferior temporal gyrus; MTG = middle temporal gyrus; Occip = occipital.

Supplementary Table 2. Fixated Word Surprisal Cluster Information, 0-225 ms

| Region | Max Cluster Duration [ms] | Max Cluster Sources | Corrected p-value |
|---|---|---|---|
| L aFus | 35 | 4 | 0.019 |
| L pFus | 23 | 3 | 0.019 |
| L aITG | 18 | 3 | 0.019 |
| L pITG | 40 | 5 | 0.019 |
| L aMTG | 25 | 5 | 0.019 |
| L pMTG | 65 | 3 | 0.019 |
| L Lateral Occip | 41 | 6 | 0.030 |
| L Parahippocampal | 37 | 16 | 0.019 |
| L Entorhinal | 46 | 6 | 0.019 |
| R pMTG | 22 | 6 | 0.035 |
| R Lingual | 70 | 30 | 0.019 |
| R Lateral Occip | 43 | 5 | 0.019 |
| R Cuneus | 35 | 3 | 0.028 |
| R Entorhinal | 33 | 5 | 0.019 |

Abbreviations: a = anterior; p = posterior; Fus = fusiform gyrus; ITG = inferior temporal gyrus; MTG = middle temporal gyrus; Occip = occipital.

Supplementary Table 3. Fixated Word Frequency in High/Low Surprisal, 0-225 ms

| | High Surprisal | | | Low Surprisal | | | |
|---|---|---|---|---|---|---|---|
| Region | Max Duration [ms] | Max Cluster Source s | Cor. p-value | Max Duration [ms] | Max Cluster Sources | Cor. p-value | ** = p < 0.05 in both Contexts |
| L aFus | 30 | 6 | .019 | 11 | 3 | .037 | ** |
| L pFus | 62 | 17 | .019 | 16 | 3 | .034 | ** |
| L aITG | 28 | 7 | .019 | 41 | 8 | .020 | ** |
| L pITG | 43 | 8 | .019 | – | – | 1.00 | high only |
| L aMTG | 25 | 5 | .020 | – | – | 1.00 | high only |
| L pMTG | 28 | 15 | .019 | 31 | 3 | 0.022 | ** |
| L Lingual | 11 | 3 | .044 | – | – | 1.00 | high only |
| L Lateral Occip | 51 | 10 | .019 | 22 | 3 | 0.019 | ** |
| L Pericalcarine | – | – | 1.00 | 23 | 6 | 0.022 | low only |
| L Cuneus | – | – | 1.00 | – | – | 1.00 | |
| L Parahippocampal | 39 | 14 | 0.019 | 14 | 13 | .019 | ** |
| L Entorhinal | – | – | 1.00 | – | – | 1.00 | |
| R aFus | – | – | 1.00 | – | – | 1.00 | |
| R pFus | – | – | 1.00 | – | – | 1.00 | |
| R aITG | – | – | 1.00 | 26 | 10 | 0.020 | low only |
| R pITG | – | – | 1.00 | – | – | 1.00 | |
| R aMTG | – | – | 1.00 | 25 | 17 | 0.019 | low only |
| R pMTG | – | – | 1.00 | 47 | 10 | 0.022 | low only |
| R Lingual | – | – | 1.00 | – | – | 1.00 | |
| R Lateral Occip | 48 | 7 | 0.019 | 10 | 3 | 0.032 | ** |
| R Pericalcarine | – | – | 1.00 | – | – | 1.00 | |
| R Cuneus | – | – | 1.00 | – | – | 1.00 | |

| | | | | | | | |
|---|---|---|---|---|---|---|---|
| R Parahippocampal | – | – | 1.00 | 33 | 8 | 0.035 | low only |
| R Entorhinal | – | – | 1.00 | 36 | 6 | 0.019 | low only |

Abbreviations: a = anterior; p = posterior; Fus = fusiform gyrus; ITG = inferior temporal gyrus; MTG = middle temporal gyrus; Occip = occipital.

Supplementary Table 4. Right Neighbour Frequency Effects, 0-225 ms

| Region | Max Cluster Duration [ms] | Max Cluster Sources | Corrected p-value |
|---|---|---|---|
| L aFus | 20 | 3 | 0.020 |
| L pFus | 39 | 6 | 0.019 |
| L Lingual | 49 | 39 | 0.019 |
| L Lateral Occip | 88 | 28 | 0.019 |
| L Pericalcarine | 54 | 4 | 0.019 |
| L Parahippocampal | 30 | 10 | 0.019 |
| L Entorhinal | 26 | 5 | 0.024 |
| R pFus | 17 | 6 | 0.041 |
| R pITG | 41 | 3 | 0.019 |
| R Lateral Occip | 31 | 11 | 0.019 |

Supplementary Table 5. Right Neighbour Surprisal Effects, 0-225 ms

| Region | Max Cluster Duration [ms] | Max Cluster Sources | Corrected p-value |
|---|---|---|---|
| L pFus | 31 | 13 | 0.019 |
| L pITG | 103 | 25 | 0.019 |
| L aMTG | 51 | 19 | 0.019 |
| L Lingual | 29 | 20 | 0.043 |
| L Lateral Occip | 66 | 8 | 0.019 |
| R pFus | 17 | 8 | 0.019 |
| R pITG | 23 | 5 | 0.037 |
| R Pericalcarine | 27 | 8 | 0.028 |
| R Cuneus | 28 | 4 | 0.019 |
| R Entorhinal | 22 | 5 | 0.032 |

Supplementary Table 6. Right Word Frequency in High/Low Right Word Surprisal, 0-225 ms
Underlined regions show main effects of Right Word Surprisal
Bold and underlined regions additionally show main effects of Right Word Frequency across all examined fixations

| | High Surprise | | | Low Surprise | | | |
|---|---|---|---|---|---|---|---|
| Region | Max Duration [ms] | Max Cluster Sources | Corrected. p-value | Max Duration [ms] | Max Cluster Sources | Corrected p-value | ** = p < 0.05 in both Contexts |
| L aFus | 31 | 8 | 0.019 | 61 | 12 | .019 | ** |
| L pFus | – | – | 1 | 42 | 9 | .019 | |
| L aITG | 18 | 3 | 0.030 | 24 | 6 | .019 | ** |
| L pITG | 35 | 3 | 0.019 | 42 | 13 | .019 | ** |
| L aMTG | 40 | 8 | 0.019 | 20 | 3 | .034 | ** |
| L pMTG | – | – | 1 | 32 | 8 | .019 | |
| **L Lingual** | 37 | 23 | 0.024 | 95 | 56 | .019 | ** |
| **L Lateral Occip** | 61 | 18 | 0.019 | 91 | 24 | .019 | ** |
| L Pericalcarine | 25 | 4 | 0.026 | 54 | 18 | .019 | ** |
| L Cuneus | – | 00 | 1 | 42 | 5 | .019 | |
| L Parahippocampal | – | – | 1 | 46 | 18 | .019 | |
| L Entorhinal | – | – | 1 | 50 | 12 | .019 | |
| R aFus | – | – | 1 | 12 | 4 | .019 | |
| **R pFus** | 41 | 15 | 0.019 | 23 | 13 | .019 | ** |
| R aITG | 21 | 9 | 0.020 | – | – | 1 | |
| R pITG | 18 | 4 | 0.032 | – | – | 1 | |
| R aMTG | 36 | 7 | 0.019 | 17 | 7 | .019 | ** |
| R pMTG | – | – | 1 | 80 | 13 | .019 | |
| R Lingual | 30 | 11 | 0.019 | 54 | 47 | .019 | ** |
| R Lateral Occip | 33 | 6 | 0.019 | 67 | 22 | .019 | ** |

| | | | | | | | |
|---|---|---|---|---|---|---|---|
| R Pericalcarine | – | – | 1 | 49 | 20 | .019 | |
| R Cuneus | – | – | 1 | 45 | 9 | .019 | |
| R Parahippocampal | 19 | 6 | 0.019 | 28 | 10 | .019 | ** |
| R Entorhinal | – | –- | 1 | 16 | 4 | .019 | ** |

Abbreviations: a = anterior; p = posterior; Fus = fusiform gyrus; ITG = inferior temporal gyrus; MTG = middle temporal gyrus; Occip = occipital. Underlined areas show main effects of right neighbour surprisal. Bold and underlined areas show both main effects of right neighbour surprisal and main effects of right neighbour frequency, across all examined fixations.

Supplementary Table 7. Right Word Frequency when next word is Skipped/Fixated, 0-225 ms

| | Next Word: Skipped | | | Next Word: Fixated | | | |
|---|---|---|---|---|---|---|---|
| Region | Max Duration [ms] | Max Cluster Source s | Corrected. p-value | Max Duration [ms] | Max Cluster Sources | Corrected p-value | Pattern |
| L aFus | 25 | 5 | 0.014* | 20 | 4 | 0.014* | Both |
| L pFus | 38 | 8 | 0.023* | | | 1 | Skipped |
| L aITG | | | 1 | 39 | 5 | 0.040* | Fixated |
| L pITG | 36 | 5 | 0.014* | 31 | 9 | 0.014* | Both |
| L aMTG | 21 | 4 | 0.032* | 12 | 9 | 0.014* | Both |
| L pMTG | 19 | 3 | 0.014* | | | 1 | Skipped |
| L Lingual | 46 | 21 | 0.014* | | | 1 | Skipped |
| L Lateral Occip | 32 | 11 | 0.014* | 16 | 3 | 0.152 | Skipped |
| L Pericalcarine | 43 | 3 | 0.014* | | | 1 | Skipped |
| L Cuneus | | | 1 | | | 1 | |
| L Parahippocampal | | | 1 | 16 | 3 | 0.014* | Fixated |
| L Entorhinal | 20 | 3 | 0.014* | 29 | 4 | 0.014* | Both |
| R aFus | | | 1 | | | 1 | |
| R pFus | 39 | 12 | 0.014* | | | 1 | Skipped |
| R aITG | 11 | 6 | 0.023* | | | 1 | Skipped |
| R pITG | | | 1 | | | 1 | |
| R aMTG | 57 | 3 | 0.014* | 15 | 8 | 0.014* | Both |
| R pMTG | 46 | 16 | 0.014* | 17 | 9 | 0.040* | Both |
| R Lingual | 18 | 4 | 0.014* | | | 1 | Skipped |
| R Lateral Occip | 33 | 7 | 0.014* | 21 | 5 | 0.014* | Both |
| R Pericalcarine | 14 | 3 | 0.014* | | | 1 | Skipped |
| R Cuneus | 11 | 4 | 0.014* | 19 | 6 | 0.014* | Both |

| | | | | | | | |
|---|---|---|---|---|---|---|---|
| R Parahippocampal | | | 1 | | | 1 | |
| R Entorhinal | | | 1 | 26 | 4 | 0.014* | Fixated |

Supplementary Table 8. Right Word Surprisal when next word is Skipped/Fixated, 0-225 ms

| | Next Word: Skipped | | | Next Word: Fixated | | | |
|---|---|---|---|---|---|---|---|
| Region | Max Duration [ms] | Max Cluster Source s | Corrected. p-value | Max Duration [ms] | Max Cluster Sources | Corrected p-value | Pattern |
| L aFus | | | 1 | | | 1 | |
| L pFus | 25 | 6 | 0.014* | | | 1 | Skipped |
| L aITG | | | 1 | | | 1 | |
| L pITG | 28 | 6 | 0.014* | 18 | 4 | 0.032* | Both |
| L aMTG | 41 | 13 | 0.023* | 16 | 3 | 0.145 | Skipped |
| L pMTG | 30 | 8 | 0.014* | | | 1 | Skipped |
| L Lingual | | | 1 | | | 1 | |
| L Lateral Occip | 36 | 6 | 0.014* | | | 1 | Skipped |
| L Pericalcarine | 16 | 3 | 0.079 | | | 1 | Skipped |
| L Cuneus | 25 | 8 | 0.023* | | | 1 | Skipped |
| L Parahippocampal | 27 | 15 | 0.032* | 12 | 11 | 0.014* | Both |
| L Entorhinal | | | 1 | | | 1 | |
| R aFus | | | 1 | 81 | 12 | 0.014* | Fixated |
| R pFus | | | 1 | 22 | 4 | 0.014* | Fixated |
| R aITG | | | 1 | 17 | 8 | 0.014* | Fixated |
| R pITG | | | 1 | 33 | 10 | 0.014* | Fixated |
| R aMTG | 35 | 12 | 0.014* | | | 1 | Skipped |
| R pMTG | 31 | 7 | 0.023* | | | 1 | Skipped |
| R Lingual | | | 1 | 11 | 3 | 0.214 | |
| R Lateral Occip | 37 | 8 | 0.014* | | | 1 | Skipped |
| R Pericalcarine | | | 1 | | | 1 | |
| R Cuneus | 23 | 4 | 0.014* | | | 1 | Skipped |

| | | | | | | | |
|---|---|---|---|---|---|---|---|
| R Parahippocampal | 11 | 8 | 0.032* | 32 | 14 | 0.023* | Both |
| R Entorhinal | 37 | 5 | 0.023* | 18 | 4 | 0.023* | Both |

Supplementary Table 9. Word Frequency in High/Low Word Surprisal, 225-500 ms

| | High Surprise | | | Low Surprise | | | |
|---|---|---|---|---|---|---|---|
| Region | Max Duration [ms] | Max Cluster Source s | Corrected. p-value | Max Duration [ms] | Max Cluster Sources | Corrected p-value | ** = p < 0.05 in only High Surprise |
| L aFus | 19 | 6 | 0.021* | | | 1 | ** |
| L pFus | 39 | 6 | 0.021* | 15 | 5 | 0.058 | ** |
| L aITG | 14 | 4 | 0.022* | 13 | 3 | 0.087 | ** |
| L pITG | | | 1 | 13 | 3 | 0.021* | |
| L aMTG | 36 | 3 | 0.021* | 28 | 5 | 0.025* | |
| L pMTG | 21 | 6 | 0.021* | | | 1 | ** |
| L Lingual | | | 1 | 16 | 3 | 0.021* | |
| L Lateral Occip | 21 | 6 | 0.021* | 29 | 5 | 0.021* | |
| L Pericalcarine | | | 1 | | | 1 | |
| L Cuneus | | | 1 | 32 | 9 | 0.021* | |
| L Parahippocampal | 22 | 7 | 0.023* | | | 1 | ** |
| L Entorhinal | 42 | 8 | 0.058 | | | 1 | |
| R aFus | 14 | 3 | 0.021* | | | 1 | ** |
| R pFus | 42 | 25 | 0.021* | 76 | 18 | 0.021* | |
| R aITG | 17 | 5 | 0.025* | | | 1 | ** |
| R pITG | 56 | 9 | 0.021* | | | 1 | ** |
| R aMTG | 13 | 3 | 0.10 | 17 | 7 | 0.021* | |
| R pMTG | 32 | 5 | 0.021* | | | 1 | ** |
| R Lingual | 19 | 20 | 0.021* | 51 | 25 | 0.021* | |
| R Lateral Occip | 76 | 23 | 0.021* | 51 | 7 | 0.021* | |
| R Pericalcarine | 14 | 3 | 0.021* | 30 | 7 | 0.021* | |
| R Cuneus | 11 | 3 | 0.059 | 31 | 4 | 0.021* | |

| | | | | | | | |
|---|---|---|---|---|---|---|---|
| R Parahippocampal | | | 1 | | | 1 | |
| R Entorhinal | 50 | 6 | 0.021* | | | 1 | ** |

Abbreviations: a = anterior; p = posterior; Fus = fusiform gyrus; ITG = inferior temporal gyrus; MTG = middle temporal gyrus; Occip = occipital. Underlined areas show main effects of right neighbour surprisal. Bold and underlined areas show both main effects of right neighbour surprisal and main effects of right neighbour frequency, across all examined fixations.

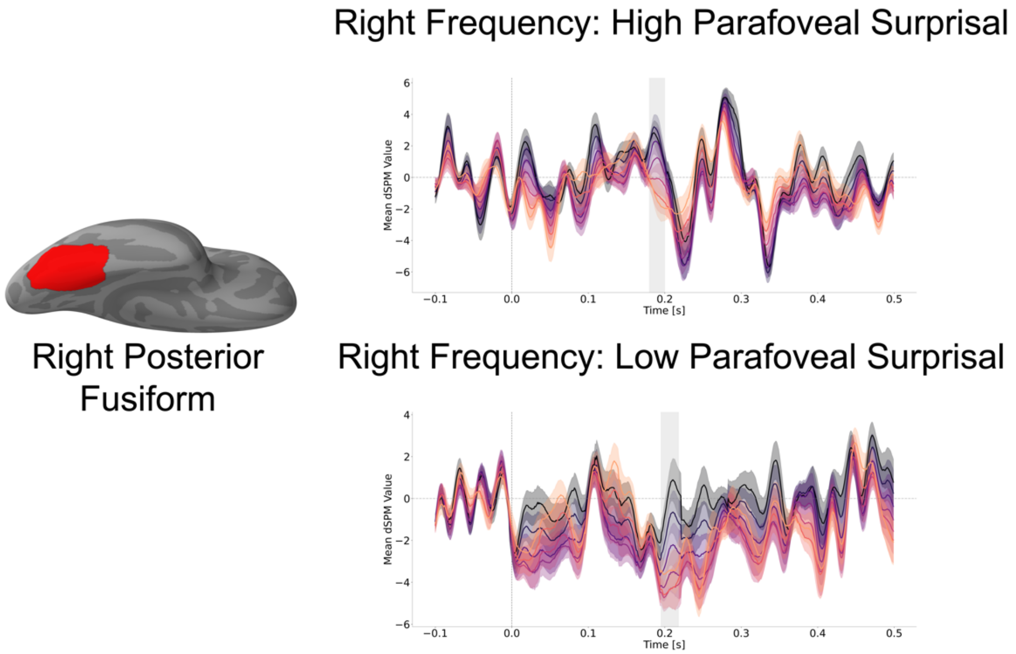


Supplementary Figure 1. Effects of frequency on high/low surprisal right neighbouring words, during the fixation on the preceding word, in the right posterior fusiform gyrus. The red shading on the ventral surface of the brain displays the ROI. Grey shading in the time course indicates the temporal extent of the significant clusters.
